# Molecular insights into electroreceptor ribbon synapses from differential gene expression in sturgeon lateral line organs

**DOI:** 10.1101/2025.02.04.636467

**Authors:** Alexander S. Campbell, Martin Minařík, David Buckley, Tanmay Anand, David Gela, Martin Pšenička, Clare V. H. Baker

## Abstract

In fishes and aquatic-stage amphibians, mechanosensory neuromasts are arranged in characteristic lines in the skin of the head and trunk, with afferent innervation from anterior or posterior lateral line nerves. In electroreceptive non-teleost jawed fishes and amphibians, fields of electrosensory ampullary organs flank some or all of the cranial neuromast lines, innervated by the anterior lateral line nerve. Like the mechanosensory hair cells found in neuromasts and the inner ear, electroreceptor cells in ampullary organs form specialised ribbon synapses with afferent nerve terminals. Ribbon synapses in hair cells are distinct from other glutamatergic synapses, including the ribbon synapses in photoreceptors: otoferlin is the Ca^2+^ sensor for synaptic vesicle exocytosis and synaptic vesicles are loaded with glutamate by vGlut3. We previously showed that the genes encoding otoferlin and vGlut3 are expressed by ampullary organs as well as neuromasts in a chondrostean ray-finned fish, the Mississippi paddlefish (*Polyodon spathula*), suggesting that electroreceptor ribbon synapses are very similar to those in hair cells. In this study, we selected seven additional synapse-related candidate genes from our previously published dataset of putatively lateral line organ-enriched genes from late-larval paddlefish, and examined their expression in developing lateral line organs in a related chondrostean, the sterlet sturgeon (*Acipenser ruthenus*). We found that genes encoding the presynaptic cell adhesion molecule Nrxn3, the calcium-independent synaptotagmin Syt14, the high-affinity glutamate re-uptake transporter EAAT1 (GLAST), calmodulin regulator protein PCP4 (PEP-19) and cell adhesion molecule DSCAML1 were expressed in both neuromasts and ampullary organs. In contrast, *Cbln18*, encoding a secreted trans-synaptic scaffolding protein, was only expressed in neuromasts and *Tulp1*, encoding tubby-related protein 1 (required for the development and function of photoreceptor ribbon synapses), was only expressed in ampullary organs. Our results support electroreceptor ribbon synapses being glutamatergic and suggest further commonalities, but also some differences, with hair cell ribbon synapses.

## Introduction

Fishes and aquatic-stage amphibians possess an evolutionarily ancient sensory system known as the lateral line (Bullock et al., 1983; Northcutt, 1997; Mogdans, 2021). Neuromasts in the skin, distributed in lines over the head and trunk, contain mechanosensory hair cells that respond to nearby water movement ("touch at a distance"), used for detecting prey or predators, obstacles, and orientation (Montgomery et al., 2014; Mogdans, 2021). Mechanical deflection of the apical ’hair bundle’ of stepped microvilli (stereocilia) leads to cation entry via the mechanoelectrical transduction complex, depolarising the cell (Cunningham and Müller, 2019; Holt et al., 2024). The hair cells in neuromasts are very similar to vestibular inner-ear hair cells (Nicolson, 2017; Shi et al., 2023). In electroreceptive non-teleost jawed fishes and amphibians, at least some neuromast lines are flanked by electrosensory ampullary organs containing electroreceptor cells (Bullock et al., 1983; Baker et al., 2013; Crampton, 2019). These have voltage-gated calcium channels in the apical membrane (identified as Cav1.3 in sharks and skates; Bellono et al., 2017; Bellono et al., 2018) that respond to low-frequency cathodal (exterior negative) environmental stimuli, such as electric fields around other animals in water, primarily for detecting prey or predators (Bullock et al., 1983; Bodznick and Montgomery, 2005; Crampton, 2019).

Afferent innervation for hair cells is provided by neurons in cranial lateral line ganglia, projecting via the anterior or posterior lateral line nerves (depending on the position of the neuromast) to the medial octavolateral nucleus of the hindbrain (Wullimann and Grothe, 2014). Electroreceptor afferents are provided exclusively by the anterior lateral line nerve, projecting centrally via its dorsal root to the dorsal octavolateral nucleus of the hindbrain (Bullock et al., 1983; Wullimann and Grothe, 2014). Both hair cells and electroreceptor cells (Jørgensen, 2005) form specialised ribbon synapses with afferent nerve terminals, characterised by an electron-dense presynaptic ’ribbon’ tethering a readily releasable pool of synaptic vesicles, enabling response to graded signals and sustained neurotransmitter release (Nicolson, 2015; Johnson et al., 2019; Moser et al., 2020).

In both hair cells and electroreceptors, depolarisation in response to stimulation opens voltage-gated calcium channels (Cav1.3 in hair cells) clustered at ribbon synapses, leading to synaptic vesicle exocytosis and release of neurotransmitter (glutamate in hair cells) (Bodznick and Montgomery, 2005; Nicolson, 2015; Johnson et al., 2019; Moser et al., 2020). Cav1.3 has also been identified as the voltage-sensing channel in the apical membrane of shark and skate electroreceptors (Bellono et al., 2017; Bellono et al., 2018; reviewed by Leitch and Julius, 2019). In hair cells, calcium entry via ribbon synapse-associated Cav1.3 channels triggers synaptic vesicle exocytosis by binding to otoferlin, a transmembrane Ca^2+^ sensor that interacts directly with Cav1.3 and membrane-fusion machinery specifically in hair cells (Hams et al.,

2017; Michalski et al., 2017; Johnson et al., 2019). Hair cell ribbon synapses are also unique in that their synaptic vesicles are filled with glutamate by the vesicular glutamate transporter vGlut3, whereas photoreceptor ribbon synapses use vGlut1 and vGlut2 (Johnson et al., 2019; Moser et al., 2020). However, very little information is available about the molecular nature of electroreceptor ribbon synapses.

We previously reported expression of *Cacna1d* (encoding the pore-forming subunit of Cav1.3), *Otof* (encoding otoferlin) and *Slc17a8* (encoding vGlut3) in late-larval ampullary organs as well as neuromasts in a chondrostean ray-finned fish, the Mississippi paddlefish (*Polyodon spathula*) (Modrell et al., 2017a). These genes were identified via a differential RNA-seq screen in late-larval paddlefish comparing the transcriptomes of operculum (rich in ampullary organs, plus some neuromasts) versus fin (similar tissue composition, but without lateral line organs) (Modrell et al., 2017a). Their conserved expression suggested that electroreceptor ribbon synapses operate in the same way as hair cell ribbon synapses. We also identified a few genes likely to be important specifically for electroreceptor function: two voltage-gated potassium channel genes (*Kcna5* and *Kcnab3*) and a beta-parvalbumin gene (encoding a calcium-buffering protein) were expressed in paddlefish ampullary organs but not neuromasts (Modrell et al., 2017a). We also identified conserved expression of many developmental genes, suggesting that electroreceptors are closely related to hair cells, as well as some developmental genes (encoding transcription factors and signalling pathway components) expressed specifically in either ampullary organs or neuromasts (Modrell et al., 2017a; Modrell et al., 2017b; Minařík et al., 2024a; Minařík et al., 2024b; Campbell et al., 2024).

Here, we report the expression in developing neuromasts and/or ampullary organs of seven synapse-related candidate genes from our paddlefish dataset of ∼500 putatively lateral line organ-enriched genes (Modrell et al., 2017a), in a more experimentally tractable chondrostean, the sterlet sturgeon (*Acipenser ruthenus*).

## Methods

### Collection, staging and fixation of sterlet yolksac larvae

Yolksac larvae of sterlet sturgeon (*Acipenser ruthenus*) were obtained at the Research Institute of Fish Culture and Hydrobiology, Faculty of Fisheries and Protection of Waters, University of South Bohemia in České Budějovice (Vodňany, Czech Republic). Stundl et al. (2022) provide detailed information about adult sterlet husbandry, *in vitro* fertilisation and raising embryos and yolk-sac larvae. Larvae were staged according to Dettlaff et al. (1993). Following euthanasia by anaesthetic overdose with MS-222 (Sigma-Aldrich), larvae were fixed for 3 hours at room temperature in modified Carnoy’s fixative (6 volumes 100% ethanol: 3 volumes 37% formaldehyde: 1 volume glacial acetic acid) and graded into ethanol before storing at -20°C.

All experimental procedures were approved by the Animal Research Committee of the Faculty of Fisheries and Protection of Waters in Vodňany, University of South Bohemia in České Budějovice, Czech Republic, and by the Ministry of Agriculture of the Czech Republic (reference number: MSMT-12550/2016-3). Experimental fish were maintained according to the principles of the European Union (EU) Harmonized Animal Welfare Act of the Czech Republic, and Principles of Laboratory Animal Care and National Laws 246/1992 “Animal Welfare” on the protection of animals.

### Gene cloning

To make late-larval sterlet cDNA, Trizol (Invitrogen, Thermo Fisher Scientific) was used to extract total RNA from stage 45 sterlet larval heads. After treating with DNAse using the Ambion Turbo DNA-free kit (Invitrogen, Thermo Fisher Scientific), cDNA was synthesised using the High-Capacity cDNA Reverse Transcription Kit (Applied Biosystems), following the manufacturers’ instructions. Genes were selected from the late-larval paddlefish (*Polyodon spathula*) lateral line organ-enriched gene-set (National Center for Biotechnology Information [NCBI] Gene Expression Omnibus (RRID:SCR_005012) accession code GSE92470; Modrell et al., 2017a). Using a Basic Local Alignment Search Tool (BLAST) database generated from our sterlet transcriptome assemblies (from pooled stages 40-45 larval sterlet heads; Minařík et al., 2024a), which are available at DDBJ/EMBL/GenBank under the accessions GKLU00000000 and GKEF01000000, the appropriate paddlefish transcriptome sequence was used in a command-line search to identify homologous sterlet sequences. NCBI BLAST (https://blast.ncbi.nlm.nih.gov/Blast.cgi; RRID:SCR_004870; McGinnis and Madden, 2004) was used to check sterlet sequence identity. PCR primers (Supplementary Table S1) were designed with Primer3Plus (RRID:SCR_003081; Untergasser et al., 2012), which is also incorporated into the Editor program of Benchling (https://benchling.com; RRID:SCR_013955). The primers were used to amplify cDNA fragments from sterlet cDNA under standard PCR conditions. After cloning the cDNA fragments into Qiagen’s pDrive cloning vector using the Qiagen PCR Cloning Kit (Qiagen), the clones were checked by sequencing (Department of Biochemistry Sequencing Facility, University of Cambridge). Sequence identity was confirmed using NCBI BLAST. For *Syt14*, *Dscaml1* and *Dscam*, synthetic gene fragments with added M13 forward and reverse primer adaptors were designed using sterlet transcriptome data and bought from Twist Bioscience.

These sterlet riboprobe template sequences were designed before the publication of chromosome-level genome assemblies for sterlet: Du et al. (2020) and the 2022 NCBI RefSeqGene (RRID:SCR_013787) reference genome assembly (GCF_902713425.1/). For all genes described here except *Cbln18*, both ohnologs (i.e., gene paralogs resulting from the whole-genome duplication) have been retained from the whole-genome duplication that occurred in the sterlet lineage (Du et al., 2020). Our phylogenetic analysis of *Cbln* family genes showed that the single *Cbln18* ohnolog in the reference sterlet genome (GCF_902713425.1) has been mis-annotated as *Cbln3* (Figure 2; Supplementary Table S1). Supplementary Table S1 includes the percentage match of each riboprobe with each ohnolog, obtained by using NCBI BLAST to perform a nucleotide BLAST search against the reference genome assembly (NCBI RefSeqGene assembly GCF_902713425.1/). The percentage match with the “targeted” ohnolog ranged from 99.3-100%. The percentage match with the second ohnolog (where present) ranged from 94.5-100%, suggesting that transcripts from the second ohnolog are also targeted by our riboprobes. Supplementary Table S1 also gives the GenBank (RRID:SCR_002760) accession numbers for each riboprobe’s top match, and the nucleotide range targeted by each riboprobe.

**Figure 1.**
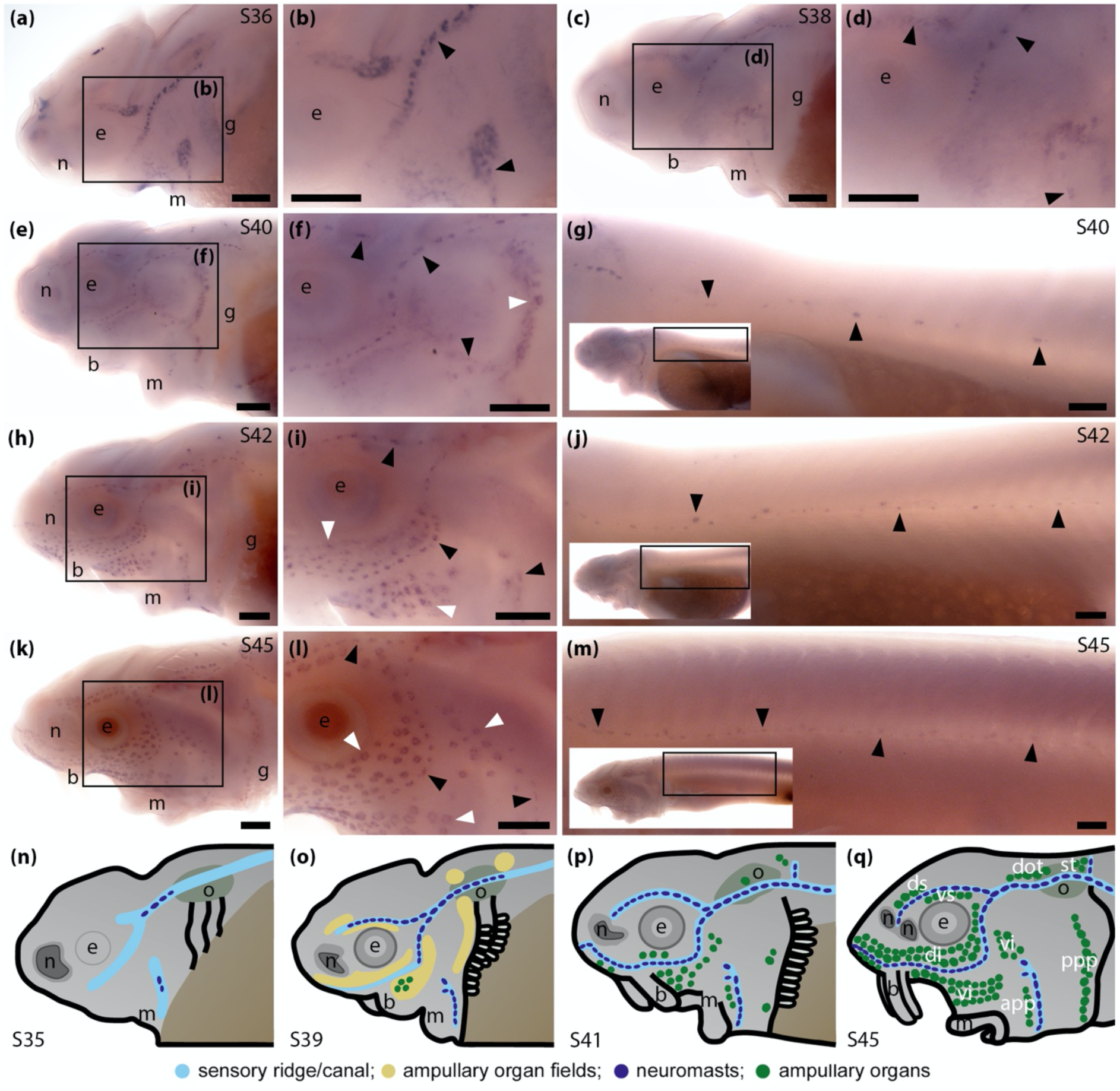
Sterlet *Nrxn3* is expressed in neuromasts and ampullary organs. *In situ* hybridisation in sterlet for *Nrxn3*. Black arrowheads indicate examples of neuromasts; white arrowheads indicate examples of ampullary organs. For images of the trunk, boxes on low-power insets delineate the regions shown. **(a-d)** *Nrxn3* expression is seen in neuromasts on the head from stage 36 (a,b), persisting at stage 38 (c,d). **(e-g)** At stage 40, *Nrxn3* expression is maintained in cranial neuromasts and is also now seen in ampullary organs (e,f) and neuromasts on the trunk (g). **(h-j)** At stage 42, *Nrxn3* expression persists in cranial neuromasts and ampullary organs (h,i) and in neuromasts on the trunk (j). **(k-m)** At stage 45, *Nrxn3* expression is maintained in neuromasts and ampullary organs on the head (k,l) and neuromasts on the trunk (m). **(n-q)** Schematic illustrations of sterlet lateral line organ development at similar stages (previously published in Minařík et al., 2024a). Abbreviations: app, anterior preopercular ampullary organ field; b, barbel; di, dorsal infraorbital ampullary organ field; dot, dorsal otic ampullary organ field; ds, dorsal supraorbital ampullary organ field; e, eye; g, gill filaments; m, mouth; n, naris; o, otic vesicle; ppp, posterior preopercular ampullary organ field; S, stage; st, supratemporal ampullary organ field; vi, ventral infraorbital ampullary organ field; vs, ventral supraorbital ampullary organ field. Scale bar: 250 μm.

**Figure 2.**
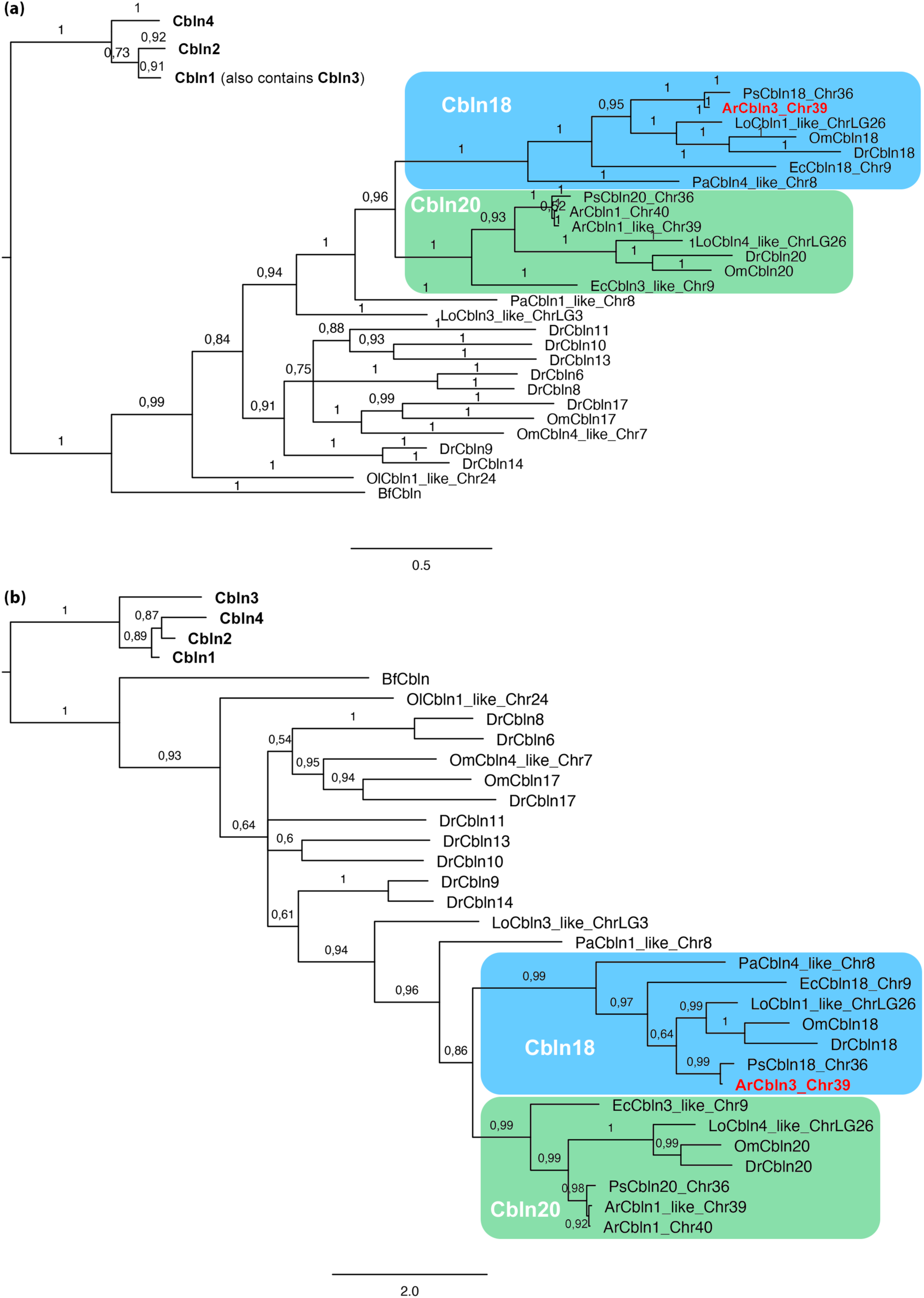
Phylogenetic analysis confirms that the sterlet ortholog of the lateral line organ-enriched paddlefish *cerebellin* transcript encodes Cbln18. **(a,b)** Selected portions of phylogenetic trees generated by Bayesian phylogenetic analysis of cerebellin family amino acid sequences using (a) MrBayes v.3.2.7 (Ronquist et al., 2012) and (b) PhyloBayes MPI v1.8c (Lartillot et al., 2013). The complete trees are provided in Supplementary Figure S1 and Supplementary Figure S2, respectively. Sequence names reflect the reference-genome annotation and show the chromosomal (Chr) location of the gene if multiple *cerebellin* genes in that species share the same or similar annotation. Maximum support for the Cbln18 clade in both trees shows that the lateral line-enriched paddlefish *cerebellin* transcript (Modrell et al., 2017) and its sterlet ortholog (highlighted in bold red font) encode Cbln18 and have been mis-annotated as *Cbln3* in the respective reference genomes (paddlefish GCF_017654505.1; sterlet GCF_902713425.1). Both trees also show that the sterlet *Cbln20* ohnologs have been mis-annotated in the reference genome as *Cbln1-like* (chromosome 39) and *Cbln1* (chromosome 40). GenBank accession numbers for the sequences used (104 from vertebrates plus the single amphioxus cerebellin sequence) are given in Supplementary Table S2. Species abbreviations: Ar, *Acipenser ruthenus*; Am, *Alligator mississippiensis*; Bf, *Branchiostoma floridae*; Cl, *Canis lupus*; Cm, *Callorhinchus milii*; Dr, *Danio rerio*; Ec, *Erpetoichthys calabaricus*; Gg, *Gallus gallus*; Hs, *Homo sapiens*; Lo, *Lepisosteus oculatus*; Mm, *Mus musculus*; Ol, *Oryzias latipes*; Om, *Oncorhynchus mykiss*; Pa, *Protopterus annectens*; Ps, *Polyodon spathula*; Sc, *Scyliorhinus canicula*; Xt, *Xenopus tropicalis*.

### Phylogenetic analysis of cerebellins

Orthologs of the target sequence (encoded by the gene annotated *Cbln3* in the sterlet reference genome assembly; GCF_902713425.1/) were checked in the SHOOT.bio database (https://shoot.bio; Emms and Kelly, 2022). The target sequence clustered with different cerebellin 18 sequences. To confirm its nature, amino acid sequences from different cerebellin families (104 vertebrate sequences plus the sole amphioxus cerebellin sequence; Supplementary Table S2) were downloaded from GenBank (RRID:SCR_002760) and aligned using the online version of MAFFT 7 (https://mafft.cbrc.jp/alignment/server/; RRID:SCR_011811; Katoh et al., 2019) with the G-INS-i iterative refinement method. Two Bayesian phylogenetic analyses were run with the aligned data matrix. First, a search in MrBayes v.3.2.7 (RRID:SCR_012067; Ronquist et al., 2012) was performed: two analyses of 10 million MC^3^ chains were run, sampling the chains every 1000 generations. These were averaged over all possible substitution models with the option ‘aamodelpr=mixed’. Trees were rooted with mid-point rooting. The second Bayesian phylogenetic analysis was run in PhyloBayes MPI v1.8c (RRID:SCR_006402; Lartillot et al., 2013) in the server CIPRES Science Gateway (https://www.phylo.org/; RRID:SCR_008439; Miller et al., 2010) under the infinite mixture CAT-GTR model. The tree was rooted with mid-point rooting. The resulting trees were visualised and edited in FigTree v1.4.4 (RRID:SCR_008515; Rambaut 2018) and Adobe Photoshop (RRID:SCR_014199; Adobe Systems Inc.).

### *In situ* hybridisation

Digoxigenin-labelled riboprobes were prepared as previously described (Minařík et al., 2024a). Wholemount *in situ* hybridisation (ISH) was performed as described in Modrell et al. (2011). At least three larvae were used per stage.

### Image capture and processing

Sterlet larvae were imaged using either a QImaging MicroPublisher 5.0 RTV camera controlled by QCapture Pro 7.0 software (RRID:SCR_014432; QImaging) or a MicroPublisher 6 color CCD camera (Teledyne Photometrics) controlled by Ocular software (RRID:SCR_024490; Teledyne Photometrics), fitted to a Leica MZFLIII dissecting microscope. Helicon Focus software (RRID:SCR_014462; Helicon Soft Limited) was used for focus stacking of image-stacks collected by focusing manually through the sample. Images were processed with Adobe Photoshop (RRID:SCR_014199; Adobe Systems Inc.).

## Results

### *Nrxn3*, encoding a presynaptic cell adhesion molecule, is expressed in sterlet neuromasts and ampullary organs

*Neurexin 3* (*Nrxn3*) was the only neurexin gene in the late-larval paddlefish lateral line organ-enriched dataset (3.1-fold enriched in operculum versus fin tissue; Modrell et al., 2017a). Neurexins are presynaptic cell adhesion molecules that function as synaptic organisers, forming trans-synaptic bridges with multiple ligands including secreted cerebellins and post-synaptic transmembrane neuroligins (reviewed by Gomez et al., 2021; Südhof, 2023). RNA-seq studies have identified *Nrxn3* expression in embryonic and post-natal hair cells in mouse (Elkon et al., 2015; Scheffer et al., 2015; Sadler et al., 2020) and zebrafish (Lush et al., 2019). A recent study confirmed expression of the long alpha form of both *nrxn3a* and *nrxn3b* in zebrafish hair cells and reported mutant analysis showing that *nrxn3* is required for the maturation of ribbon synapses in zebrafish neuromasts, and in the inner ear of zebrafish and mouse (Jukic et al., 2024).

We performed wholemount *in situ* hybridisation (ISH) for sterlet *Nrxn3* at a range of larval stages starting from stage 36 (hatching; 6 days post-fertilisation; Zeiske et al., 2003). This is shortly after the first *Cacna1d*-positive differentiated hair cells are seen in cranial neuromasts at stage 35 (Minařík et al., 2024a). The first differentiated electroreceptors (identified by *Cacna1d* or *Kcnab3* expression) are not seen until a few days later, at stages 40-41 (Minařík et al., 2024a). At stage 36, *Nrxn3* was expressed in neuromasts on the head (Figure 1a,b). Expression in cranial neuromasts was maintained at stage 38 (Figure 1c,d) and at stage 40, when expression was also seen in some ampullary organs and in neuromasts on the trunk (Figure 1e-g). *Nrxn3* expression persisted in neuromasts and ampullary organs at stage 42 (Figure 1h-j) and at stage 45 (Figure 1k-m; the onset of independent feeding; 14 days post-fertilisation; Zeiske et al., 2003). Figure 1n-q shows schematics (previously published in Minařík et al., 2024a) illustrating the progression of cranial neuromast and ampullary organ development in sterlet. The spatiotemporal pattern of sterlet *Nrxn3* expression correlates with the appearance of differentiated hair cells and electroreceptors (Minařík et al., 2024a), suggesting it is most likely expressed in electroreceptors as well as hair cells. Given that Nrxn3 is required for the maturation of ribbon synapses in zebrafish and mouse hair cells (Jukic et al., 2024), our sterlet data suggest that Nrxn3 may also be important for the maturation of electroreceptor ribbon synapses.

### *Cbln18*, encoding a secreted trans-synaptic scaffolding glycoprotein, is expressed in sterlet neuromasts but not ampullary organs

*Cerebellin 18* (*Cbln18*) was one of the most highly lateral line organ-enriched genes (and the only cerebellin gene) identified in our late-larval paddlefish differential RNA-seq screen (33.3-fold enriched in operculum versus fin tissue; Modrell et al., 2017a). Cerebellins are secreted scaffolding glycoproteins that bridge synapses by forming tripartite complexes with presynaptic neurexins (specifically, the long alpha forms of Nrxn1-3) and post-synaptic proteins, such as the delta glutamate receptors GluD1 and GluD2 (for Cbln1-3) or the netrin receptors neogenin-1 and DCC (for Cbln4) (Ferrer-Ferrer and Dityatev, 2018; Südhof, 2023).

Phylogenetic analysis of vertebrate cerebellin proteins confimed the identity of the lateral line organ-enriched paddlefish *cerebellin* transcript (Modrell et al., 2017a) and its sterlet ortholog as *Cbln18* (Figure 2a,b; Supplementary Figures S1, S2; Supplementary Table S2). The sterlet genome underwent a lineage-specific whole-genome duplication (Du et al., 2020) but only a single *Cbln18* ohnolog (on chromosome 39) was identified in the reference genome (GCF_902713425.1), where it has been misannotated as *Cbln3* (Figure 2; Supplementary Figures S1,S2). The paddlefish *Cbln18* gene has also been misannotated as *Cbln3* in the reference genome (GCF_017654505.1) (Figure 2a,b; Supplementary Figures S1, S2). However, *Cbln3* seems to be mammal-specific: no Cbln proteins from non-mammalian vertebrates cluster with Cbln3 (Supplementary Figures S1,S2).

No information has been published about the expression or function of zebrafish *cbln18*. ISH in sterlet showed strong *Cbln18* expression in neuromasts on the head at stage 36 and stage 38 (Figure 3a-d) and on both the trunk and head from stage 40 until stage 45 (the latest stage examined) (Figure 3e-m). No expression was detected at any stage in ampullary organs (in which electroreceptors start to differentiate from stages 40-41; Minařík et al., 2024a). This suggests that Cbln18 is important specifically for the synapses between hair cells and their afferent axon terminals, but not for electroreceptor synapses.

**Figure 3.**
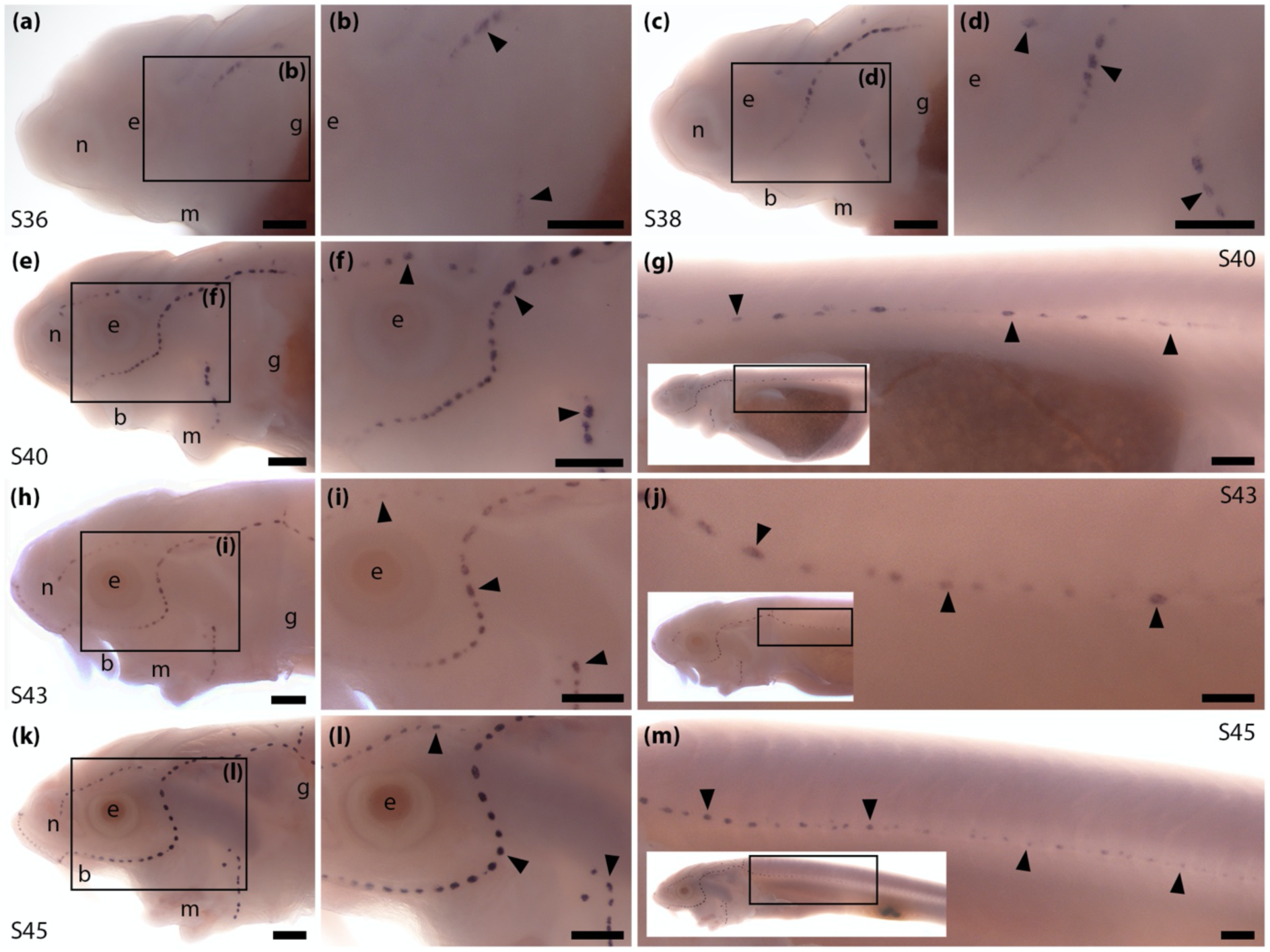
Sterlet *Cbln18* is expressed in neuromasts but not ampullary organs. *In situ* hybridisation in sterlet for *Cbln18*. Black arrowheads indicate examples of neuromasts. For images of the trunk, boxes on low-power insets delineate the regions shown. **(a-d)** *Cbln18* expression is seen in neuromasts on the head at stage 36 (a,b), persisting at stage 38 (c,d). **(e-g)** At stage 40, expression is maintained in cranial neuromasts (e,f) and is also visible in trunk neuromasts (g). (**h-m**) At stages 42 (h-j) and 45 (k-m), *Cbln18* expression persists in neuromasts on the head (h,i,k,l) and trunk (j,m). Abbreviations: b, barbel; e, eye; g, gill filaments; m, mouth; n, naris; S, stage. Scale bar: 250 μm.

### *Slc1a3*, encoding excitatory amino acid transporter 1 (EAAT1), is expressed in sterlet neuromasts and ampullary organs

*Solute carrier family 1 member 3* (*Slc1a3*) was 11.3-fold lateral line organ-enriched in late-larval paddlefish (Modrell et al., 2017a). *Slc1a3* encodes excitatory amino acid transporter 1 (EAAT1), also known as glial high-affinity glutamate transporter and glutamate/aspartate transporter (GLAST). In the CNS, EAAT1 is found in the membrane of astrocytes where it acts to clear glutamate from the synaptic cleft after synaptic transmission (reviewed by Niciu et al., 2012; Andersen et al., 2021). In the rodent inner ear, EAAT1 is expressed at the basolateral membranes of cochlear and vestibular supporting cells (Furness and Lehre, 1997; Takumi et al., 1997) and required for glutamate uptake from cochlear hair cell ribbon synapses (Glowatzki et al., 2006; Chen et al., 2010). Zebrafish *slc1a3* is also expressed in neuromasts (Gesemann et al., 2010).

Preliminary ISH data from late-larval paddlefish (not shown) identified the expression of *Slc1a3* in both neuromasts and ampullary organs. We undertook a more detailed study in sterlet. *Slc1a3* was not detectably expressed by ISH at stage 36 (when some differentiated neuromast hair cells are already present; Minařík et al., 2024a) or stage 38 (Figure 4a,b). However, by stage 40 faint expression was seen in cranial neuromasts (Figure 4c,d). At stage 42, this expression had extended to ampullary organs (Figure 4e,f) and trunk neuromasts (Figure 4g). At stage 45, strong expression was seen in ampullary organs and weaker expression in neuromasts on both head and trunk (Figure 4h-j). Neuromast expression was in a ring-like pattern (compare *Slc1a3* expression in Figure 4i with *Nrxn3* expression in Figure 1l), suggesting *Slc1a3* is expressed by supporting cells. These data suggest that EAAT1 (GLAST) expressed by supporting cells may clear glutamate from the ribbon synapses of both hair cells and electroreceptors.

**Figure 4.**
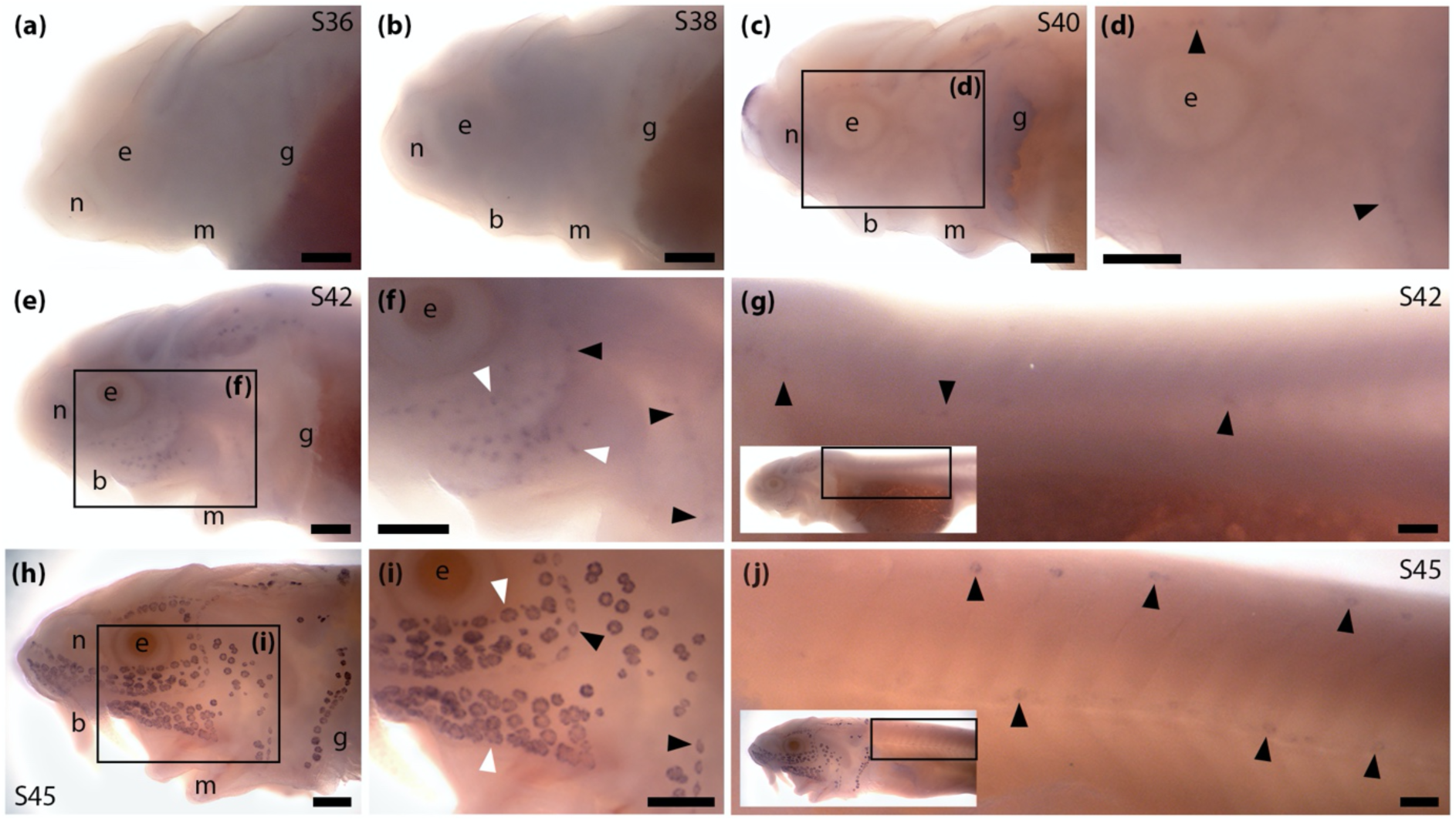
Sterlet *Slc1a3* (encoding EAAT1/GLAST) is expressed in neuromasts and ampullary organs. *In situ* hybridisation in sterlet for *Slc1a3*. Black arrowheads indicate examples of neuromasts; white arrowheads indicate examples of ampullary organs. For images of the trunk, boxes on low-power insets delineate the location of the trunk regions shown. **(a,b)** *Slc1a3* expression is not seen at stage 36 (a) or stage 38 (b). **(c,d)** At stage 40, *Slc1a3* expression is observed in cranial neuromasts. **(e-g)** At stage 42, *Slc1a3* expression persists in cranial neuromasts and is also seen in ampullary organs (e,f) and trunk neuromasts (g). **(h-j)** At stage 45, *Slc1a3* expression persists on the head in ampullary organs and neuromasts (h,i) and more weakly in trunk neuromasts. The ring-like expression pattern in neuromasts (seen most clearly in cranial neuromasts, panel i) suggests *Slc1a3* is expressed by supporting cells rather than hair cells (compare with *Nrxn3* expression in neuromasts in Figure 1l). Abbreviations: b, barbel; e, eye; g, gill filaments; m, mouth; n, naris; S, stage. Scale bar: 250 μm.

### *Pcp4*, encoding calmodulin-regulator protein PCP4 (PEP-19), is expressed in sterlet neuromasts and ampullary organs

*Purkinje cell protein 4* (*Pcp4*), was 13.2-fold lateral line organ-enriched in late-larval paddlefish (Modrell et al., 2017a). *Pcp4* encodes calmodulin-regulator protein PCP4,Pcp4 (also known as PEP-19), which binds to the intracellular Ca^2+^ sensor calmodulin in a Ca^2+^-independent manner and can suppress calmodulin-dependent signalling, preventing cellular Ca^2+^ overload and protecting against glutamate-induced excitotoxicity in neurons (Slemmon et al., 1996; Slemmon et al., 2000; Kanazawa et al., 2008; Wang et al., 2013). *Pcp4* is expressed in mouse inner-ear hair cells (Thomas et al., 2003; Burns et al., 2015; Cai et al., 2015) but its role in hair cells is unknown.

Preliminary ISH data from late-larval paddlefish (not shown) suggested *Pcp4* was expressed in both neuromasts and ampullary organs. In sterlet, ISH revealed strong *Pcp4* expression in neuromasts on the head already at stage 36 (shortly after the first differentiated hair cells are seen at stage 35; Minařík et al., 2024a) and stage 38 (Figure 5a-d), and on the head and trunk at stage 40, stage 42 and stage 45 (Figure 5e-m). At stage 45, weaker *Pcp4* expression was also seen in ampullary organs (Figure 5k,l), suggesting PCP4 is important for the function of electroreceptors, as well as hair cells. However, differentiated electroreceptors are first seen from stages 40-41 (Minařík et al., 2024a). The lack of detectable *Pcp4* expression at stage 42, versus strong *Pcp4* expression in neuromasts at stage 36, shortly after the first differentiated hair cells are seen (Minařík et al., 2024a), suggests that the function of PCP4 may differ in ampullary organs and neuromasts.

**Figure 5.**
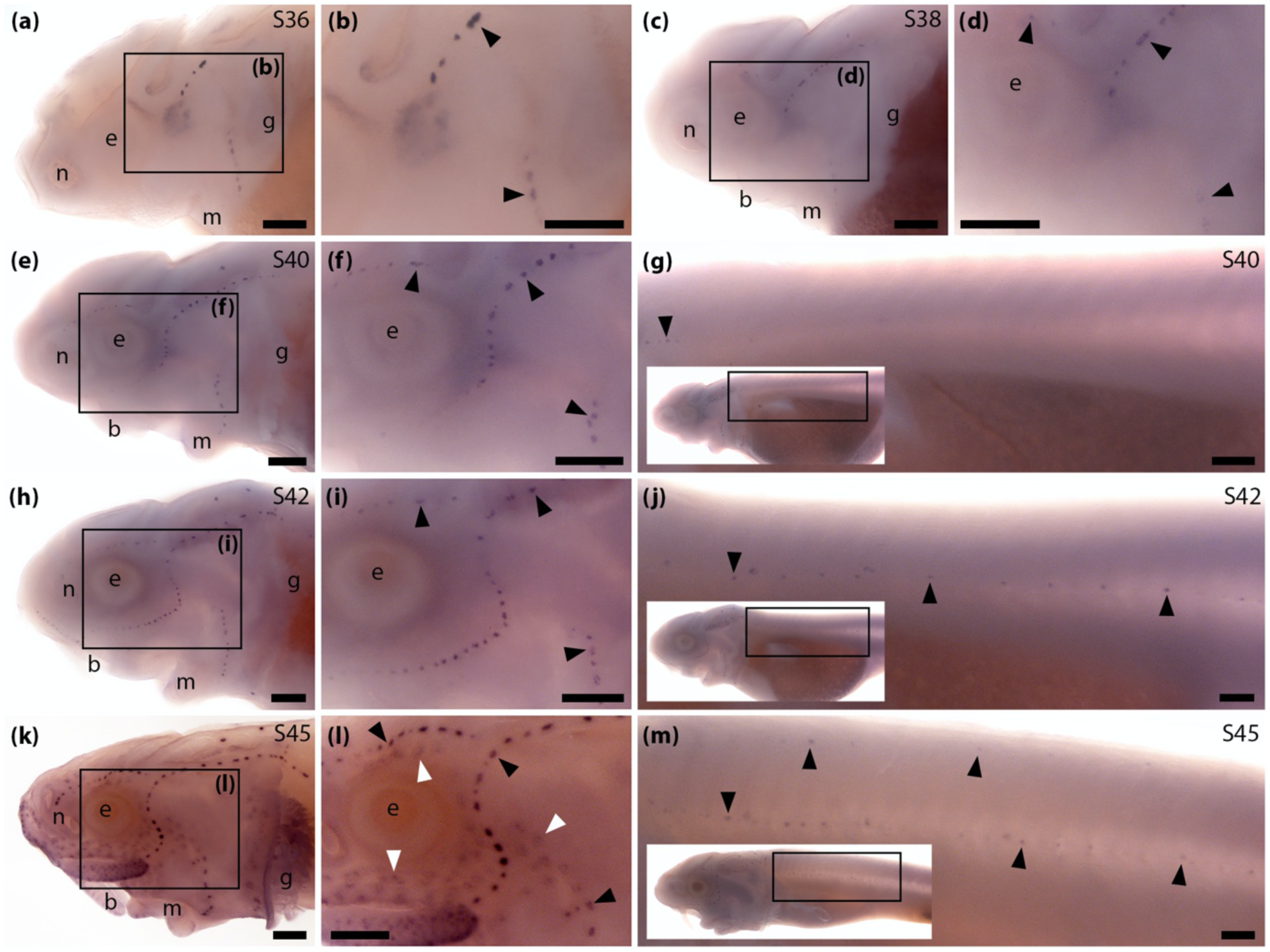
Sterlet *Pcp4* is expressed in neuromasts and ampullary organs. *In situ* hybridisation in sterlet for *Pcp4*. Black arrowheads indicate examples of neuromasts; white arrowheads indicate examples of ampullary organs. For images of the trunk, boxes on low-power insets delineate the location of the trunk regions shown. **(a-d)** *Pcp4* expression is seen in neuromasts on the head at stage 36 (a,b) and stage 38 (c,d). **(e-j)** At stage 40 (e-g) and stage 42 (h-j), *Pcp4* expression is maintained in neuromasts on the head and is also visible in neuromasts on the trunk. (**k-m**) At stage 45, neuromast expression is maintained on the head and weaker expression in ampullary organs also appears (k,l). On the trunk, expression continues in neuromasts (m). Abbreviations: b, barbel; e, eye; g, gill filaments; m, mouth; n, naris; S, stage. Scale bar: 250 μm.

### *Syt14*, encoding a calcium-independent synaptotagmin, is expressed in sterlet neuromasts and ampullary organs

*Synaptotagmin 14* (*Syt14*) was 12.0-fold lateral line organ-enriched in late-larval paddlefish (Modrell et al., 2017a). Synaptotagmins are transmembrane proteins with two cytoplasmic Ca^2+^-binding C2 domains (reviewed by Wolfes and Dean, 2020). *Syt14* is expressed in embryonic and postnatal inner ear hair cells in mouse (Scheffer et al., 2015) and in utricular (vestibular) hair cells in chicken (Scheibinger et al., 2022). Many synaptotagmins are important for the regulation of calcium-dependent membrane fusion events, with some acting as fast calcium sensors for synaptic vesicle exocytosis (Wolfes and Dean, 2020). In contrast, Syt14 is calcium-independent and its precise function is unknown (Fukuda, 2003; Wolfes and Dean, 2020).

In sterlet, ISH for *Syt14* revealed weak expression in developing neuromasts at stage 38 and stage 40 (Figure 6a-f) and in ampullary organs at stage 42 (Figure 6g,h). At stage 45, *Syt14* expression was seen in ampullary organs and, more weakly, in neuromasts on both head and trunk (Figure 6i-k). These data suggest that the (unknown) function of Syt14 in hair cells is likely conserved in electroreceptors.

**Figure 6.**
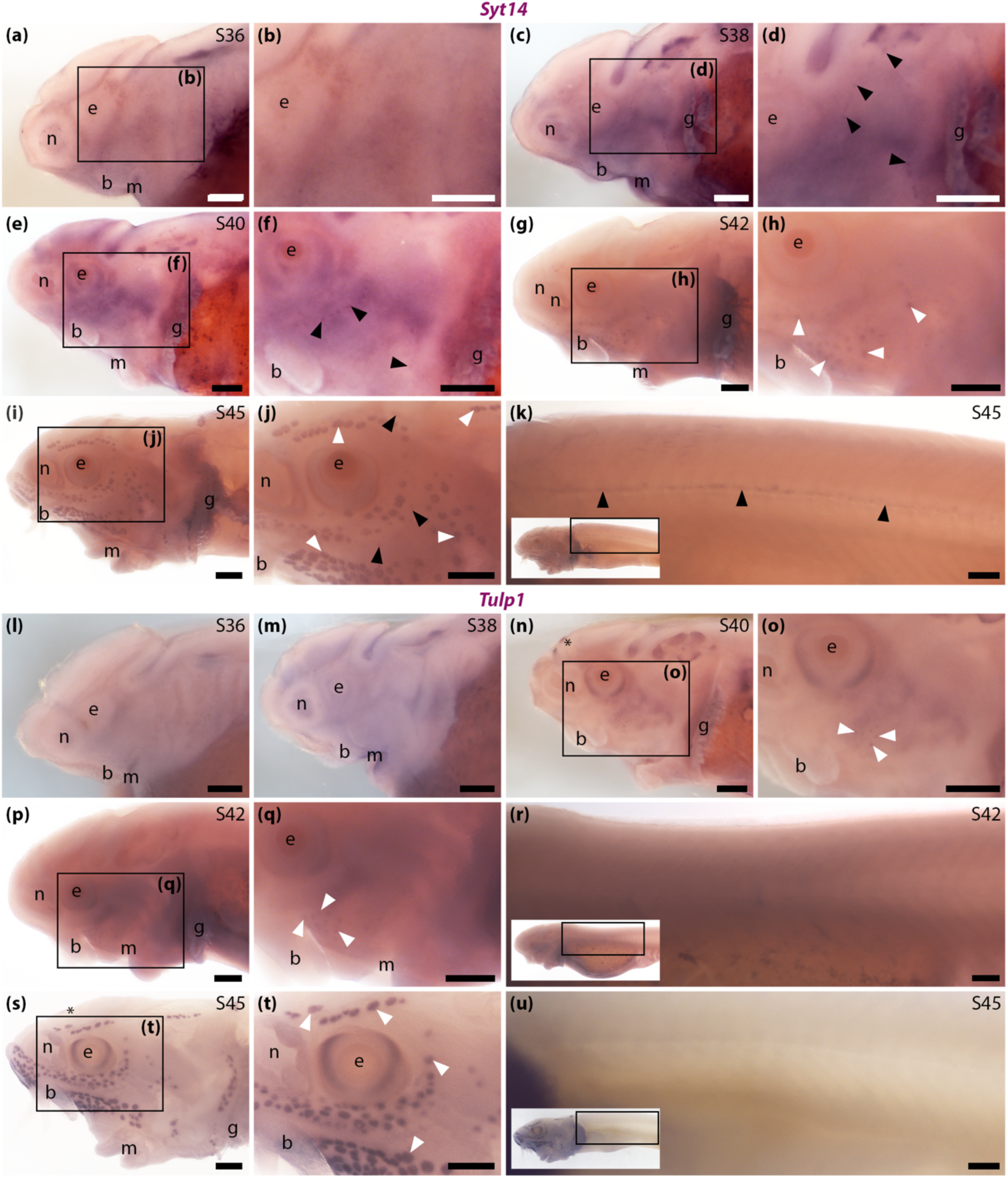
Sterlet neuromasts and ampullary organs express *Syt14*, but only ampullary organs express *Tulp1*. *In situ* hybridisation in sterlet. Black arrowheads indicate examples of neuromasts; white arrowheads indicate examples of ampullary organs. For images of the trunk, boxes on low-power insets delineate the location of the trunk regions shown. **(a,b)** No specific *Syt14* expression is seen at stage 36. **(c-f)** At stage 38 (c,d) and stage 40 (e,f), weak *Syt14* expression is observed in cranial neuromasts. **(g,h)** At stage 42, weak *Syt14* expression is seen in ampullary organs. **(i-k)** At stage 45, *Syt14* expression is maintained in ampullary organs (h,i) and weaker expression is seen in neuromasts on the head (i,j and trunk (k). **(l,m)** No specific *Tulp1* expression is seen at either stage 36 (l) or stage 38 (m). **(n,o)** At stage 40, *Tulp1* expression is observed in the eye, a midline patch between the eyes that is most likely the epiphysis (asterisk), and weakly in ampullary organs. No expression is seen in neuromasts. (**p-r**) At stage 42, *Tulp1* expression continues in ampullary organs but not in neuromasts, either on the head (p,q) or trunk (r). **(s-u)** At stage 45, *Tulp1* expression is maintained in the eye, weakly in the presumed epiphysis (asterisk), and in ampullary organs (s,t). No expression is seen in neuromasts, either on the head (s,t) or trunk (u). Abbreviations: b, barbel; e, eye; g, gill filaments; m, mouth; n, naris; S, stage. Scale bar: 250 μm.

### *Tulp1*, encoding tubby-related protein 1, is expressed in sterlet ampullary organs but not neuromasts

*TUB like protein 1* (*Tulp1*) was 2.4-fold lateral line organ-enriched in late-larval paddlefish (Modrell et al., 2017a). Tulp1 is a member of the tubby family of proteins, characterised by a conserved C-terminal "tubby" domain that binds phosphoinositide 4,5-bisphosphate (PIP2), linking the protein to the membrane (Mukhopadhyay and Jackson, 2011). In mice, Tulp1 is restricted to the retina, where it interacts with Ribeye (the main structural component of the presynaptic ribbon) and is required for the normal development and function of the photoreceptor ribbon synapse (Grossman et al., 2009; Wahl et al., 2016).

ISH showed no detectable expression of sterlet *Tulp1* at stage 36 or stage 38 (Figure 6l,m). At stage 40 and stage 42, *Tulp1* expression was seen in the eye (presumably in the retina) and, weakly, in ampullary organs, but not in neuromasts (Figure 6n-r). At stage 45, *Tulp1* expression was maintained in the eye and in ampullary organs (Figure 6s,t). No *Tulp1* expression was seen in neuromasts, either on the head (Figure 6s,t) or trunk (Figure 6u). Thus, Tulp1 seems to be ampullary organ-specific within the developing lateral line system. Given the importance of Tulp1 for the development and function of the photoreceptor ribbon synapse (Grossman et al., 2009; Wahl et al., 2016), these data suggest that Tulp1 may be important specifically for the ribbon synapse of electroreceptors, but not hair cells.

### *Dscaml1*, encoding cell adhesion molecule DSCAML1, is expressed in sterlet neuromasts and ampullary organs

*Down’s syndrome cell adhesion molecule-like 1* (*Dscaml1*) was 25.3-fold lateral line organ-enriched in late-larval paddlefish (Modrell et al., 2017a). *Dscaml1* encodes cell adhesion molecule DSCAML1, a homophilic transmembrane cell adhesion molecule of the immunoglobulin superfamily (Agarwala et al., 2001). In the mouse retina, *Dscaml1* is expressed in rod photoreceptors, rod bipolar cells (which also form ribbon synapses, like photoreceptors) and AII amacrine cells (Fuerst et al., 2009). In *Dscaml1*-null retinas, ribbon synapses between rod bipolar cells and AII amacrine cells are functional but seem to be immature, with fewer ribbons, some floating (detached from the presynaptic membrane), significantly more synaptic vesicles and a slower decay of the synaptic current (Fuerst et al., 2009; also see Garrett et al., 2016).

In sterlet, ISH for *Dscaml1* revealed strong expression in neuromasts on the head at stage 36 and stage 38 (Figure 7a-d), and on the head and trunk at stage 40, stage 42 and stage 45 (Figure 7e-m), although at stage 45, expression seemed to be fading. At stage 40, stage 42 and stage 45, *Dscaml1* expression was also seen in ampullary organs (Figure 7e-m).

**Figure 7.**
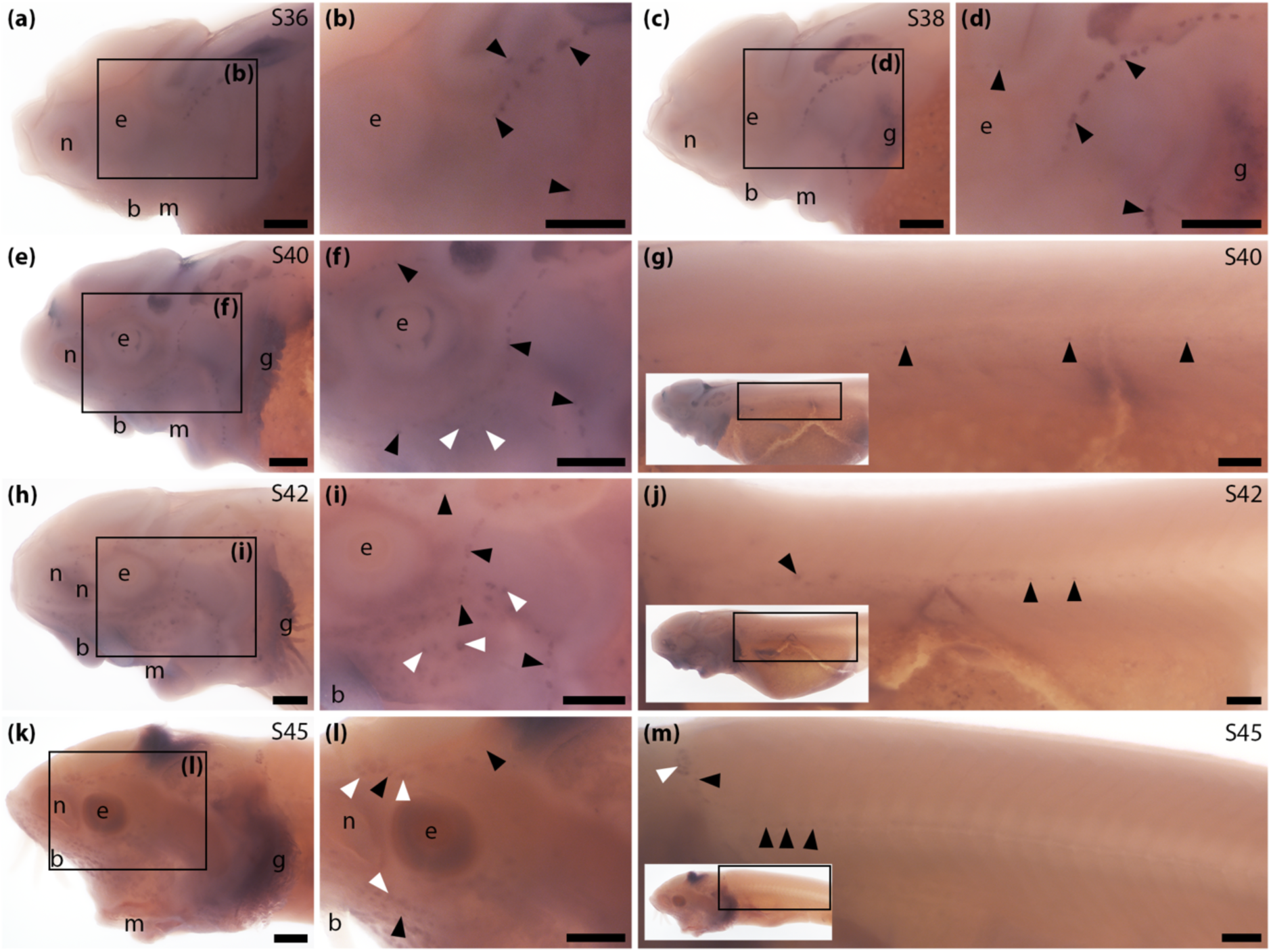
Sterlet *Dscaml1* is expressed in neuromasts and ampullary organs. *In situ* hybridisation in sterlet for *Dscaml1*. Black arrowheads indicate examples of neuromasts; white arrowheads indicate examples of ampullary organs. For images of the trunk, boxes on low-power insets delineate the location of the trunk regions shown. **(a-d)** *Dscaml1*expression is seen in neuromasts on the head at stage 36 (a,b) and stage 38 (c,d). **(e-j)** At stage 40 (e-g) and stage 42 (h-j), *Dscaml1* expression continues in neuromasts on the head and is also seen in neuromasts on the trunk, as well as ampullary organs. (**k-m**) At stage 45, *Dscaml1* expression is maintained in ampullary organs (k,l) but neuromast expression seems weaker, especially on the trunk where it is only detectable rostrally (m). Abbreviations: b, barbel; e, eye; g, gill filaments; m, mouth; n, naris; S, stage. Scale bar: 250 μm.

We also tested expression in sterlet of the related gene *Dscam*, which was 3.5-fold enriched in late-larval paddlefish (Modrell et al., 2017a). However, no specific *Dscam* expression was seen in developing lateral line organs (data not shown).

## Discussion

We studied the developmental expression of seven synapse-related genes selected from a paddlefish lateral line organ-enriched transcriptome (Modrell et al., 2017a), in a related chondrostean fish, the sterlet sturgeon. Their various expression patterns provide insights into the structure and potential physiology of ribbon synapses in electroreceptors, identifying both conserved aspects and molecular differences with ribbon synapses in hair cells.

### Expression of *Slc1a3*, encoding EAAT1 (GLAST), supports electroreceptor synapses being glutamatergic

Hair cell ribbon synapses are glutamatergic (see Johnson et al., 2019; Moser et al., 2020), but the excitatory neurotransmitter at electroreceptor synapses has not been identified (see Bodznick and Montgomery, 2005; Leitch and Julius, 2019). We previously reported the expression in late-larval paddlefish neuromasts and ampullary organs of *Slc17a8*, encoding vGlut3 (Modrell et al., 2017a), which is required to refill synaptic vesicles at hair cell (but not photoreceptor) ribbon synapses (see Johnson et al., 2019; Moser et al., 2020). Here, we found that larval sterlet neuromasts and ampullary organs also express *Slc1a3*, encoding EAAT1 (GLAST). This transporter is required for supporting cells of the mammalian inner ear to take up glutamate from hair cell ribbon synapses (Glowatzki et al., 2006; Chen et al., 2010). Shared expression of both *Slc1a3* (EAAT1/GLAST) and *Slc17a8* (vGlut3; Modrell et al., 2017a) in ampullary organs and neuromasts strengthens the proposal that ribbon synapses are glutamatergic in electroreceptors as well as hair cells.

### Expression of *Nrxn3* and *Cbln18* sheds light on the structure of ribbon synapses in hair cells versus electroreceptors

The presynaptic cell adhesion molecule Nrxn3 was recently shown to be required for the maturation of hair-cell ribbon synapses in zebrafish and mouse (Jukic et al., 2024). We identified shared expression of *Nrxn3* in sterlet neuromasts and ampullary organs, suggesting that Nrxn3 may play the same role at electroreceptor ribbon synapses. The specific binding partners of Nrxn3 in hair cells and electroreceptors are unknown. Neurexins commonly partner directly with neuroligins in the post-synaptic membrane: for example, in the mouse cochlea, Nrxn1 and Nlgn3 are both important for the maturation of hair-cell ribbon synapses (Ramirez et al., 2022). The paddlefish lateral line organ-enriched geneset (Modrell et al., 2017a) did not include any neuroligin genes, but this is not surprising as the mRNA would be expressed in afferent lateral line neurons. However, our gene expression data suggest that one synaptic partner for Nrxn3 in sterlet hair cells, but not electroreceptors, is likely to be Cbln18, a member of the cerebellin family of secreted scaffolding glycoproteins (Ferrer-Ferrer and Dityatev, 2018; Südhof, 2023). Our phylogenetic analysis showed that *Cbln18* is one of a group of cerebellin genes not found in mammals. *Cbln18* was strongly expressed in sterlet neuromasts throughout development, but not seen at any stage in ampullary organs. In hair cells, Nrxn3 and Cbln18 likely form a tripartite complex with glutamate receptors on the post-synaptic membrane (Ferrer-Ferrer and Dityatev, 2018; Gomez et al., 2021; Südhof, 2023). *Cbln18* was the only cerebellin gene in the paddlefish lateral line organ-enriched gene-set (Modrell et al., 2017a), but this dataset is not exhaustive and it is possible that a different cerebellin gene is expressed in electroreceptors. Taken together, our data suggest that ribbon synapses in hair cells and electroreceptors include presynaptic Nrxn3, but only hair-cell ribbon synapses include secreted Cbln18.

### Expression of *Syt14*, *Tulp1* and *Dscaml1* gives further insight into ribbon synapses in hair cells versus electroreceptors

Another synapse-associated gene with shared expression in developing sterlet neuromasts and ampullary organs was *Syt14*, encoding a calcium-independent member of the synaptotagmin family (reviewed by Wolfes and Dean, 2020). The better studied family members Syt1, Syt2, Syt7 and Syt9 are transmembrane synaptic vesicle proteins that act as calcium sensors and regulate synaptic vesicle exocytosis at synapses in the central nervous system (Wolfes and Dean, 2020). However, at mature hair cell ribbon synapses, otoferlin is the calcium sensor that controls synaptic vesicle exocytosis, rather than synaptotagmins (reviewed by Johnson et al., 2019; Moser et al., 2020). Furthermore, Syt14 is calcium-independent (Fukuda, 2003). Syt14 is expressed in cerebellar Purkinje cells and human *SYT14* mutation is associated with spinocerebellar ataxia (Doi et al., 2011), but its precise function is unknown. Another calcium-independent synaptotagmin, Syt4, both inhibits vesicle exocytosis and slows endocytosis (reviewed by Wolfes and Dean, 2020). Given that we observed *Syt14* expression in both ampullary organs and neuromasts, it is possible that Syt14 could similarly be involved in regulating exocytosis and/or endocytosis at both electroreceptor and hair cell ribbon synapses.

*Dscaml1*, encoding the homophilic cell adhesion molecule DSCAML1 (Agarwala et al., 2001), was also expressed in developing neuromasts and ampullary organs in sterlet. (The related gene *Dscam* did not show specific expression in developing lateral line organs.) Data from *Dscaml1*-null mouse retinas suggested that *Dscaml1* may be required for the final maturation of ribbon synapses between Dscaml1-expressing rod bipolar cells and AII amacrine cells (Fuerst et al., 2009; Garrett et al., 2016). Although these ribbon synapses were functional in *Dscaml1*-null retinas, fewer ribbons formed; some ribbons were not associated with the presynaptic membrane (’floating’) and significantly more synaptic vesicles were present, suggesting a potential defect in exocytosis (Fuerst et al., 2009). These features were described as suggestive of immature synapses (Garrett et al., 2016). *Dscaml1* expression has not been reported in hair cells, to our knowledge; in zebrafish, *Dscaml1* expression has only been examined in the retina (Galicia et al., 2018; Ma et al., 2020). It is possible that Dscaml1 might play a similar role in the maturation of ribbon synapses between neuromast hair cells/electroreceptors and their afferent neurons, assuming the latter also express *Dscaml1*.

In contrast to the shared expression in neuromasts and ampullary organs of *Syt14* and *Dscaml1*, we saw differential expression of a gene that is important for regulating endocytosis at the photoreceptor ribbon synapse, *Tulp1* (Wahl et al., 2016). This gene, which encodes tubby-related protein 1, is mutated in a subset of human patients with retinitis pigmentosa, in which photoreceptors degenerate (reviewed by Frederick and Zenisek, 2023). In mice, Tulp1 is restricted to the retina, where it is required for endocytosis in the periactive zone of the photoreceptor ribbon synapse (Wahl et al., 2016). Such endocytosis is essential for the continued release of synaptic vesicles at the photoreceptor synapse (Wen et al., 2018). Sterlet *Tulp1* was expressed in developing ampullary organs, but not neuromasts. Expression was also seen in a dorsal midline patch likely to be the epiphysis (pineal gland), as reported for both *tulp1a* and *tulp1b* in zebrafish (Jia et al., 2022). Pinealocytes also have ribbon synapses (Moser et al., 2020). Hence, it is possible that Tulp1 could be important for vesicle recycling at the ribbon synapse of electroreceptors and pinealocytes, as it is in photoreceptor ribbon synapses (reviewed by Frederick and Zenisek, 2023).

*Tulp1* is also required for the normal development of photoreceptor ribbon synapses (Grossman et al., 2009). Tulp1 in photoreceptors interacts with Ribeye (Ebke et al., 2021), the major constituent of the presynaptic ribbon in all ribbon synapses, including in hair cells (Moser et al., 2020; Voorn and Vogl, 2020) and most likely also electroreceptors, given that paddlefish ampullary organs express the Ribeye-specific A domain of *Ctbp2*, encoding Ribeye (Modrell et al., 2017a). In *Tulp1*-null mice, few intact ribbons are found: the ribbon synapse proteins Piccolo and Bassoon fail to associate properly and the dendrites of associated bipolar cells are much shorter than normal, resulting in malformed synapses (Grossman et al., 2009). By analogy, Tulp1 may play multiple roles at the electroreceptor ribbon synapse, ranging from development to regulating endocytosis.

Tulp1 also has non-synapse-related roles in photoreceptors. In the inner segment of photoreceptors, Tulp1 binds to Kif3, a subunit of the Kinesin-2 motor complex involved in ciliary membrane protein trafficking (Ebke et al., 2021). In *tulp1*-mutant zebrafish, the primary cilium of photoreceptors is significantly shorter than normal and opsins are mis-localised (Jia et al., 2022). Electroreceptors also have an apical primary cilium (see Jørgensen, 2005; Baker and Modrell, 2018). Thus, it is possible that Tulp1 in electroreceptors also has non-synapse related roles in trafficking proteins in the ciliary membrane.

### Differential onset and levels of *Pcp4* expression suggest potential differences in its role in electroreceptors versus hair cells

Calmodulin-regulator protein PCP4 (PEP-19) binds calmodulin in a Ca^2+^-independent manner through its IQ domain (Slemmon et al., 1996). Calmodulin is an EF-hand intracellular Ca^2+^ sensor that interacts with and modifies the activity of hundreds of target proteins required for a plethora of cellular functions (Chin and Means, 2000; Berchtold and Villalobo, 2014; Hussey et al., 2023). PCP4 can downregulate Ca^2+^-calmodulin signalling and protect against glutamate-mediated excitotoxicity arising from raised levels of intracellular Ca^2+^ (Slemmon et al., 2000; Kanazawa et al., 2008; Wang et al., 2013). Thus, the expression of *Pcp4* that we identified in larval sterlet neuromasts and ampullary organs (also reported in mouse inner-ear hair cells; Thomas et al., 2003; Burns et al., 2015; Cai et al., 2015) could protect both hair cells and electroreceptors against glutamate-mediated excitotoxicity.

However, differences in the onset and relative levels of *Pcp4* expression in neuromasts versus ampullary organs may suggest potential differences in the role(s) of Pcp4 in hair cells versus electroreceptors. In neuromasts, *Pcp4* was strongly expressed in a spatiotemporal pattern correlating closely with hair cell differentiation (as previously detected by expression of *Cacna1d*, encoding the pore-forming subunit of Cav1.3; Minařík et al., 2024a). In contrast, the onset of detectable *Pcp4* expression in ampullary organs was much later relative to electroreceptor differentiation (as previously detected by the expression of *Cacna1d* or electroreceptor-specific *Kcna5*; Minařík et al., 2024a). *Pcp4* expression in ampullary organs was also weaker than in neuromasts, as seen by qualitatively comparing the intensity of ISH staining between ampullary organs and neuromasts in the same larvae. It is possible that ampullary organs use a different calmodulin regulator to protect against glutamate-mediated excitotoxicity and/or that the requirement for PCP4 differs in electroreceptors and hair cells.

One possibility for differing PCP4 requirements in electroreceptors and hair cells may be the importance of calmodulin for the rapid Ca^2+^-dependent inactivation of L-type voltage-gated calcium channels, such as Cav1.3 (Peterson et al., 1999; Zühlke et al., 1999; reviewed by Ames, 2021). After Cav1.3 opens and intracellular Ca^2+^ levels increase, Ca^2+^-bound calmodulin binds to Cav1.3, inactivating the channel (Ames, 2021). In hair cells, ribbon synapse-associated Cav1.3 channels open in response to hair cell depolarisation and the Ca^2+^ influx triggers synaptic vesicle exocytosis (Johnson et al., 2019; Moser et al., 2020). In electroreceptors, Cav1.3 channels most likely have the same function at the ribbon synapse, given that *Cacna1d* was the only pore-forming Cav channel gene identified in the paddlefish lateral line organ-enriched gene-set (Modrell et al., 2017a) and the predominant pore-forming Cav channel mRNA in skate and shark ampullary organs (Bellono et al., 2017; Bellono et al., 2018). However, Cav1.3 plays an additional role in electroreceptors: it was identified as the voltage-sensing calcium channel in the apical membrane of skate and shark electroreceptors (Bellono et al., 2017; Bellono et al., 2018; reviewed by Leitch and Julius, 2019). Electroreceptor function relies on constant activation of calcium-driven voltage oscillations mediated by low-threshold Cav1.3 channels in the apical membrane: the calcium influx in turn activates an outward potassium current (mediated by different potassium channels in skate and shark) (Bellono et al., 2017; Bellono et al., 2018; reviewed by Leitch and Julius, 2019). It is possible that the differences in timing and expression levels of the calmodulin-regulator gene *Pcp4* in sterlet ampullary organs versus neuromasts relate somehow to this additional role of Cav1.3 in the apical membrane of electroreceptors and the importance of Ca^2+^-calmodulin for rapid Ca^2+^-dependent inactivation of Cav1.3 (reviewed by Ames, 2021).

## Conclusion

We found that seven genes involved in synapse development, stability and/or function were also expressed in developing sterlet lateral line organs. The gene encoding the pre-synaptic organiser protein Nrxn3 was expressed in neuromasts and ampullary organs, as was a calcium-independent synaptotagmin gene, *Syt14*, and *Dscaml1*, encoding a homophilic cell adhesion molecule that seems to be important for the final maturation of ribbon synapses in rod bipolar cells (Fuerst et al., 2009; Garrett et al., 2016). Nrxn3 likely forms a synaptic complex with secreted Cbln18 in hair cell ribbon synapses, as *Cbln18* was expressed in neuromasts but not ampullary organs. Conversely, *Tulp1*, which is required for the development and function of ribbon synapses in photoreceptors (Grossman et al., 2009; Wahl et al., 2016), was expressed in ampullary organs but not neuromasts, suggesting Tulp1 may play similar roles in electroreceptors. Expression in sterlet neuromasts and ampullary organs of *Slc1a3*, encoding the glutamate re-uptake transporter EAAT1 (GLAST), supports ribbon synapses being glutamatergic in electroreceptors, as well as hair cells. Finally, *Pcp4* (encoding calmodulin regulator protein PCP4) was also expressed in sterlet neuromasts and ampullary organs. However, the relatively low expression level of *Pcp4* in ampullary organs versus neuromasts may suggest potential differences in PCP4 regulation of Ca^2+^-calmodulin in electroreceptors versus hair cells. Overall, our gene expression analysis has given novel molecular insights into similarities and differences between ribbon synapses in electroreceptors and hair cells.

## Acknowledgements

Thanks to Marek Rodina and Martin Kahanec for their help with sterlet spawns, and to Michaela Vazačová for her help with embryo incubation and fixation.

## Author contributions

A.S.C. led the project, performed most of the experiments, prepared most of the manuscript figures, and wrote the first draft of the manuscript. M.M. performed some of the experiments, prepared two of the manuscript figures and finalised the phylogenetic tree figures. D.B. undertook the phylogenetic analysis and provided the phylogenetic trees. T.A. performed some of the experiments and contributed to data interpretation. M.P. and D.G. were instrumental in enabling all work with sterlet embryos. C.V.H.B. conceived the project, provided guidance and helped to write the manuscript together with A.S.C. All authors read and commented on the manuscript.

## Data availability statement

The publication and associated supplementary figures include representative example images of larvae from each experiment. Additional data underlying this publication consist of further images of these and other larvae from each experiment. Public sharing of these images is not cost-efficient, but they are available from the corresponding author upon reasonable request. Previously published sterlet transcriptome assemblies (from pooled stage 40-45 sterlet heads; Minařík et al., 2024a) are available at DDBJ/EMBL/GenBank under the accessions GKLU00000000 (https://www.ncbi.nlm.nih.gov/nuccore/GKLU00000000) and GKEF01000000 (https://www.ncbi.nlm.nih.gov/nuccore/GKEF00000000.1). Previously published paddlefish RNA-seq data (from pooled paddlefish opercula and fin tissue at stage 46; Modrell et al., 2017a) are available via the NCBI Gene Expression Omnibus (GEO) database (https://www.ncbi.nlm.nih.gov/geo/) under accession code GSE92470.

## Ethics statement

Sterlet animal work was reviewed and approved by The Animal Research Committee of Research Institute of Fish Culture and Hydrobiology, Faculty of Fisheries and Protection of Waters, University of South Bohemia in České Budějovice, Vodňany, Czech Republic and Ministry of Agriculture of the Czech Republic (MSMT-12550/2016-3). Experimental fish were maintained according to the principles of the European Union (EU) Harmonized Animal Welfare Act of the Czech Republic, and Principles of Laboratory Animal Care and National Laws 246/1992 “Animal Welfare” on the protection of animals.

## Rights Retention Statement

This work was funded by a grant from the Biotechnology and Biological Sciences Research Council (BBSRC: BB/P001947/1). For the purpose of open access, the author has applied a Creative Commons Attribution (CC BY) licence to any Author Accepted Manuscript version arising.

## References

Agarwala, K.L., Ganesh, S., Tsutsumi, Y., Suzuki, T., Amano, K. & Yamakawa, K. (2001) Cloning and functional characterization of DSCAML1, a novel DSCAM-like cell adhesion molecule that mediates homophilic intercellular adhesion. Biochem. Biophys. Res. Commun., 285, 760–772.

Ames, J.B. (2021) L-Type Ca^2+^ channel regulation by calmodulin and CaBP1. Biomolecules, 11, 1811.

Andersen, J.V., Markussen, K.H., Jakobsen, E. et al. (2021) Glutamate metabolism and recycling at the excitatory synapse in health and neurodegeneration. Neuropharmacology, 196, 108719.

Baker, C.V.H. & Modrell, M.S. (2018) Insights into electroreceptor development and evolution from molecular comparisons with hair cells. Integr. Comp. Biol., 58, 329–340.

Baker, C.V.H., Modrell, M.S. & Gillis, J.A. (2013) The evolution and development of vertebrate lateral line electroreceptors. J. Exp. Biol., 216, 2515–2522.

Bellono, N.W., Leitch, D.B. & Julius, D. (2017) Molecular basis of ancestral vertebrate electroreception. Nature, 543, 391–396.

Bellono, N.W., Leitch, D.B. & Julius, D. (2018) Molecular tuning of electroreception in sharks and skates. Nature, 558, 122–126.

Berchtold, M.W. & Villalobo, A. (2014) The many faces of calmodulin in cell proliferation, programmed cell death, autophagy, and cancer. Biochim. Biophys. Acta, 1843, 398–435.

Bodznick, D. & Montgomery, J.C. (2005) The physiology of low-frequency electrosensory systems. In Electroreception, (Eds, Bullock, T.H., Hopkins, C.D., Popper, A.N. & Fay, R.R.) Springer, New York, pp. 132–153.

Bullock, T.H., Bodznick, D.A. & Northcutt, R.G. (1983) The phylogenetic distribution of electroreception: evidence for convergent evolution of a primitive vertebrate sense modality. Brain Res. Rev., 287, 25–46.

Burns, J.C., Kelly, M.C., Hoa, M., Morell, R.J. & Kelley, M.W. (2015) Single-cell RNA-Seq resolves cellular complexity in sensory organs from the neonatal inner ear. Nat. Commun., 6, 8557.

Cai, T., Jen, H.-I., Kang, H., Klisch, T.J., Zoghbi, H.Y. & Groves, A.K. (2015) Characterization of the transcriptome of nascent hair cells and identification of direct targets of the Atoh1 transcription factor. J. Neurosci., 35, 5870–5883.

Campbell, A.S., Minařík, M., Franěk, R. et al. (2024) Two opposing roles for Bmp signalling in the development of electrosensory lateral line organs. bioRxiv, doi: 10.1101/2024.03.07.583945.

Chen, Z., Kujawa, S.G. & Sewell, W.F. (2010) Functional roles of high-affinity glutamate transporters in cochlear afferent synaptic transmission in the mouse. J. Neurophysiol., 103, 2581–2586.

Chin, D. & Means, A.R. (2000) Calmodulin: a prototypical calcium sensor. Trends Cell Biol., 10, 322–328.

Crampton, W.G.R. (2019) Electroreception, electrogenesis and electric signal evolution. J. Fish Biol., 95, 92–134.

Cunningham, C.L. & Müller, U. (2019) Molecular structure of the hair cell mechanoelectrical transduction complex. Cold Spring Harb. Perspect. Med., 9, a033167.

Dettlaff, T.A., Ginsburg, A.S. & Schmalhausen, O.I. (1993) Sturgeon Fishes: Developmental Biology and Aquaculture. Springer-Verlag, Berlin.

Doi, H., Yoshida, K., Yasuda, T. et al. (2011) Exome sequencing reveals a homozygous *SYT14* mutation in adult-onset, autosomal-recessive spinocerebellar ataxia with psychomotor retardation. Am. J. Hum. Genet., 89, 320–327.

Du, K., Stöck, M., Kneitz, S. et al. (2020) The sterlet sturgeon genome sequence and the mechanisms of segmental rediploidization. *Nat*. Ecol. Evol., 4, 841–852.

Ebke, L.A., Sinha, S., Pauer, G.J.T. & Hagstrom, S.A. (2021) Photoreceptor compartment-specific TULP1 interactomes. Int. J. Mol. Sci., 22, 8066.

Elkon, R., Milon, B., Morrison, L. et al. (2015) RFX transcription factors are essential for hearing in mice. Nat. Commun., 6, 8549.

Emms, D.M. & Kelly, S. (2022) SHOOT: phylogenetic gene search and ortholog inference. Genome Biol., 23, 85.

Ferrer-Ferrer, M. & Dityatev, A. (2018) Shaping synapses by the neural extracellular matrix. Front. Neuroanat., 12, 40.

Frederick, C.E. & Zenisek, D. (2023) Ribbon synapses and retinal disease: Review. Int. J. Mol. Sci., 24, 5090.

Fuerst, P.G., Bruce, F., Tian, M. et al. (2009) DSCAM and DSCAML1 function in self-avoidance in multiple cell types in the developing mouse retina. Neuron, 64, 484–497.

Fukuda, M. (2003) Molecular cloning, expression, and characterization of a novel class of synaptotagmin (Syt XIV) conserved from *Drosophila* to humans. J. Biochem., 133, 641–649.

Furness, D.N. & Lehre, K.P. (1997) Immunocytochemical localization of a high-affinity glutamate-aspartate transporter, GLAST, in the rat and guinea-pig cochlea. Eur. J. Neurosci., 9, 1961–1969.

Galicia, C.A., Sukeena, J.M., Stenkamp, D.L. & Fuerst, P.G. (2018) Expression patterns of*dscam* and *sdk* gene paralogs in developing zebrafish retina. Mol. Vis., 24, 443–458.

Garrett, A.M., Tadenev, A.L., Hammond, Y.T., Fuerst, P.G. & Burgess, R.W. (2016) Replacing the PDZ-interacting C-termini of DSCAM and DSCAML1 with epitope tags causes different phenotypic severity in different cell populations. eLife, 5, e16144.

Gesemann, M., Lesslauer, A., Maurer, C.M., Schönthaler, H.B. & Neuhauss, S.C. (2010) Phylogenetic analysis of the vertebrate excitatory/neutral amino acid transporter (SLC1/EAAT) family reveals lineage specific subfamilies. BMC Evol. Biol., 10, 117.

Glowatzki, E., Cheng, N., Hiel, H. et al. (2006) The glutamate-aspartate transporter GLAST mediates glutamate uptake at inner hair cell afferent synapses in the mammalian cochlea. J. Neurosci., 26, 7659–7664.

Gomez, A.M., Traunmüller, L. & Scheiffele, P. (2021) Neurexins: molecular codes for shaping neuronal synapses. Nat. Rev. Neurosci., 22, 137–151.

Grossman, G.H., Pauer, G.J., Narendra, U., Peachey, N.S. & Hagstrom, S.A. (2009) Early synaptic defects in tulp1-/- mice. Invest. Ophthalmol. Vis. Sci., 50, 3074–3083.

Hams, N., Padmanarayana, M., Qiu, W. & Johnson, C.P. (2017) Otoferlin is a multivalent calcium-sensitive scaffold linking SNAREs and calcium channels. Proc. Natl. Acad. Sci. U.S.A., 114, 8023–8028.

Holt, J.R., Fettiplace, R. & Müller, U. (2024) Sensory transduction in auditory hair cells-PIEZOs can’t touch this. J. Gen. Physiol., 156, e202413585.

Hussey, J.W., Limpitikul, W.B. & Dick, I.E. (2023) Calmodulin mutations in human disease. Channels, 17, 2165278.

Jia, D., Gao, P., Lv, Y. et al. (2022) Tulp1 deficiency causes early-onset retinal degeneration through affecting ciliogenesis and activating ferroptosis in zebrafish. Cell Death Dis., 13, 962.

Johnson, S.L., Safieddine, S., Mustapha, M. & Marcotti, W. (2019) Hair cell afferent synapses: function and dysfunction. Cold Spring Harb. Perspect. Med., 9, a033175.

Jørgensen, J.M. (2005) Morphology of electroreceptive sensory organs. In Electroreception, (Eds, Bullock, T.H., Hopkins, C.D., Popper, A.N. & Fay, R.R.) Springer, New York, pp. 47–67.

Jukic, A., Lei, Z., Cebul, E.R. et al. (2024) Presynaptic Nrxn3 is essential for ribbon-synapse maturation in hair cells. Development, 151, dev202723.

Kanazawa, Y., Makino, M., Morishima, Y., Yamada, K., Nabeshima, T. & Shirasaki, Y. (2008) Degradation of PEP-19, a calmodulin-binding protein, by calpain is implicated in neuronal cell death induced by intracellular Ca2+ overload. Neuroscience, 154, 473–481.

Katoh, K., Rozewicki, J. & Yamada, K.D. (2019) MAFFT online service: multiple sequence alignment, interactive sequence choice and visualization. Brief. Bioinform., 20, 1160–1166.

Lartillot, N., Rodrigue, N., Stubbs, D. & Richer, J. (2013) PhyloBayes MPI: phylogenetic reconstruction with infinite mixtures of profiles in a parallel environment. Syst. Biol., 62, 611–615.

Leitch, D.B. & Julius, D. (2019) Electrosensory transduction: comparisons across structure, afferent response properties, and cellular physiology. In Electroreception: Fundamental Insights from Comparative Approaches, (Eds, Carlson, B.A., Sisneros, J.A., Popper, A.N. & Fay, R.R.) Springer, Cham, pp. 63–90.

Lush, M.E., Diaz, D.C., Koenecke, N. et al. (2019) scRNA-Seq reveals distinct stem cell populations that drive hair cell regeneration after loss of Fgf and Notch signaling. eLife, 8, e44431.

Ma, M., Ramirez, A.D., Wang, T., et al. (2020) Zebrafish *dscaml1* deficiency impairs retinal patterning and oculomotor function. J. Neurosci., 40, 143–158.

McGinnis, S. & Madden, T.L. (2004) BLAST: at the core of a powerful and diverse set of sequence analysis tools. Nucleic Acids Res., 32, W20–5.

Michalski, N., Goutman, J.D., Auclair, S.M. et al. (2017) Otoferlin acts as a Ca(2+) sensor for vesicle fusion and vesicle pool replenishment at auditory hair cell ribbon synapses. eLife, 6, e31013.

Miller, M.A., Pfeiffer, W. & Schwartz, T. (2010) “Creating the CIPRES Science Gateway for inference of large phylogenetic trees.” 2010 Gateway Computing Environments Workshop (GCE), New Orleans, LA, USA, 2010, 1–8.

Minařík, M., Modrell, M.S., Gillis, J.A. et al. (2024a) Identification of multiple transcription factor genes potentially involved in the development of electrosensory *versus* mechanosensory lateral line organs. Front. Cell Dev. Biol., 12, 1327924.

Minařík, M., Campbell, A.S., Franěk, R. et al. (2024b) Atoh1 is required for the formation of lateral line electroreceptors and hair cells, whereas Foxg1 represses an electrosensory fate. bioRxiv, doi: 10.1101/2023.04.15.537030.

Modrell, M.S., Bemis, W.E., Northcutt, R.G., Davis, M.C. & Baker, C.V.H. (2011) Electrosensory ampullary organs are derived from lateral line placodes in bony fishes. Nat. Commun., 2, 496.

Modrell, M.S., Lyne, M., Carr, A.R. et al. (2017a) Insights into electrosensory organ development, physiology and evolution from a lateral line-enriched transcriptome. eLife, 6, e24197.

Modrell, M.S., Tidswell, O.R.A. & Baker, C.V.H. (2017b) Notch and Fgf signaling during electrosensory versus mechanosensory lateral line organ development in a non-teleost ray-finned fish. Dev. Biol., 431, 48–58.

Mogdans, J. (2021) Physiology of the peripheral lateral line system. In The Senses: A Comprehensive Reference, (Ed, Fritzsch, B.) Elsevier, pp. 143–162.

Montgomery, J., Bleckmann, H. & Coombs, S. (2014) Sensory ecology and neuroethology of the lateral line. In The Lateral Line System, (Eds, Coombs, S.C., Bleckmann, H., Fay, R.R. & Popper, A.N.) Springer, New York, pp. 121–150.

Moser, T., Grabner, C.P. & Schmitz, F. (2020) Sensory processing at ribbon synapses in the retina and the cochlea. Physiol. Rev., 100, 103–144.

Mukhopadhyay, S. & Jackson, P.K. (2011) The tubby family proteins. Genome Biol., 12, 225.

Niciu, M.J., Kelmendi, B. & Sanacora, G. (2012) Overview of glutamatergic neurotransmission in the nervous system. Pharmacol. Biochem. Behav., 100, 656–664.

Nicolson, T. (2015) Ribbon synapses in zebrafish hair cells. Hear. Res., 330, 170–177.

Nicolson, T. (2017) The genetics of hair-cell function in zebrafish. J. Neurogenet., 31, 102–112.

Northcutt, R.G. (1997) Evolution of gnathostome lateral line ontogenies. Brain Behav. Evol.,50, 25–37.

Peterson, B.Z., DeMaria, C.D., Adelman, J.P. & Yue, D.T. (1999) Calmodulin is the Ca^2+^ sensor for Ca^2+^ -dependent inactivation of L-type calcium channels. Neuron, 22, 549–558.

Rambaut, A. (2018) FigTree–Tree Figure Drawing Tool Version v. 1.4. 4. Institute of Evolutionary Biology, *University of Edinburgh*: *Edinburgh*.

Ramirez, M.A., Ninoyu, Y., Miller, C. et al. (2022) Cochlear ribbon synapse maturation requires Nlgn1 and Nlgn3. iScience, 25, 104803.

Ronquist, F., Teslenko, M., van der Mark, P. et al. (2012) MrBayes 3.2: efficient Bayesian phylogenetic inference and model choice across a large model space. Syst. Biol., 61, 539–542.

Sadler, E., Ryals, M.M., May, L.A. et al. (2020) Cell-specific transcriptional responses to heat shock in the mouse utricle epithelium. Front. Cell. Neurosci., 14, 123.

Scheffer, D.I., Shen, J., Corey, D.P. & Chen, Z.-Y. (2015) Gene expression by mouse inner ear hair cells during development. J. Neurosci., 35, 6366–6380.

Scheibinger, M., Janesick, A., Benkafadar, N., Ellwanger, D.C., Jan, T.A. & Heller, S. (2022) Cell-type identity of the avian utricle. Cell Rep., 40, 111432.

Shi, T., Beaulieu, M.O., Saunders, L.M. et al. (2023) Single-cell transcriptomic profiling of the zebrafish inner ear reveals molecularly distinct hair cell and supporting cell subtypes. eLife, 12, e82978.

Slemmon, J.R., Feng, B. & Erhardt, J.A. (2000) Small proteins that modulate calmodulin-dependent signal transduction: effects of PEP-19, neuromodulin, and neurogranin on enzyme activation and cellular homeostasis. Mol. Neurobiol., 22, 99–113.

Slemmon, J.R., Morgan, J.I., Fullerton, S.M., Danho, W., Hilbush, B.S. & Wengenack, T.M. (1996) Camstatins are peptide antagonists of calmodulin based upon a conserved structural motif in PEP-19, neurogranin, and neuromodulin. J. Biol. Chem., 271, 15911–15917.

Stundl, J., Soukup, V., Franěk, R. et al. (2022) Efficient CRISPR mutagenesis in sturgeon demonstrates its utility in large, slow-maturing vertebrates. Front. Cell Dev. Biol., 10, 750833.

Südhof, T.C. (2023) Cerebellin–neurexin complexes instructing synapse properties. Curr. Opin. Neurobiol., 81, 102727.

Takumi, Y., Matsubara, A., Danbolt, N.C. et al. (1997) Discrete cellular and subcellular localization of glutamine synthetase and the glutamate transporter GLAST in the rat vestibular end organ. Neuroscience, 79, 1137–1144.

Thomas, S., Thiery, E., Aflalo, R. et al. (2003) PCP4 is highly expressed in ectoderm and particularly in neuroectoderm derivatives during mouse embryogenesis. Gene Expr. Patterns, 3, 93–97.

Untergasser, A., Cutcutache, I., Koressaar, T. et al. (2012) Primer3 - new capabilities and interfaces. Nucl. Acids Res., 40, e115.

Voorn, R.A. & Vogl, C. (2020) Molecular assembly and structural plasticity of sensory ribbon synapses-a presynaptic perspective. Int. J. Mol. Sci., 21, 8758.

Wahl, S., Magupalli, V.G., Dembla, M. et al. (2016) The disease protein Tulp1 is essential for periactive zone endocytosis in photoreceptor ribbon synapses. J. Neurosci., 36, 2473–2493.

Wang, X., Xiong, L.W., El Ayadi, A., Boehning, D. & Putkey, J.A. (2013) The calmodulin regulator protein, PEP-19, sensitizes ATP-induced Ca2+ release. J. Biol. Chem., 288, 2040–2048.

Wen, X., Van Hook, M.J., Grassmeyer, J.J. et al. (2018) Endocytosis sustains release at photoreceptor ribbon synapses by restoring fusion competence. J. Gen. Physiol., 150, 591–611.

Wolfes, A.C. & Dean, C. (2020) The diversity of synaptotagmin isoforms. Curr. Opin. Neurobiol., 63, 198–209.

Wullimann, M.F. & Grothe, B. (2014) The central nervous organization of the lateral line system. In The Lateral Line System, (Eds, Coombs, S.C., Bleckmann, H., Fay, R.R. & Popper, A.N.) Springer, New York, pp. 195–251.

Zeiske, E., Kasumyan, A., Bartsch, P. & Hansen, A. (2003) Early development of the olfactory organ in sturgeons of the genus *Acipenser*: a comparative and electron microscopic study. Anat. Embryol., 206, 357–372.

Zühlke, R.D., Pitt, G.S., Deisseroth, K., Tsien, R.W. & Reuter, H. (1999) Calmodulin supports both inactivation and facilitation of L-type calcium channels. Nature, 399, 159–162.

