## Supplementary Figures S1 and S2 for "Molecular insights into electroreceptor ribbon synapses from differential gene expression in sturgeon lateral line organs"

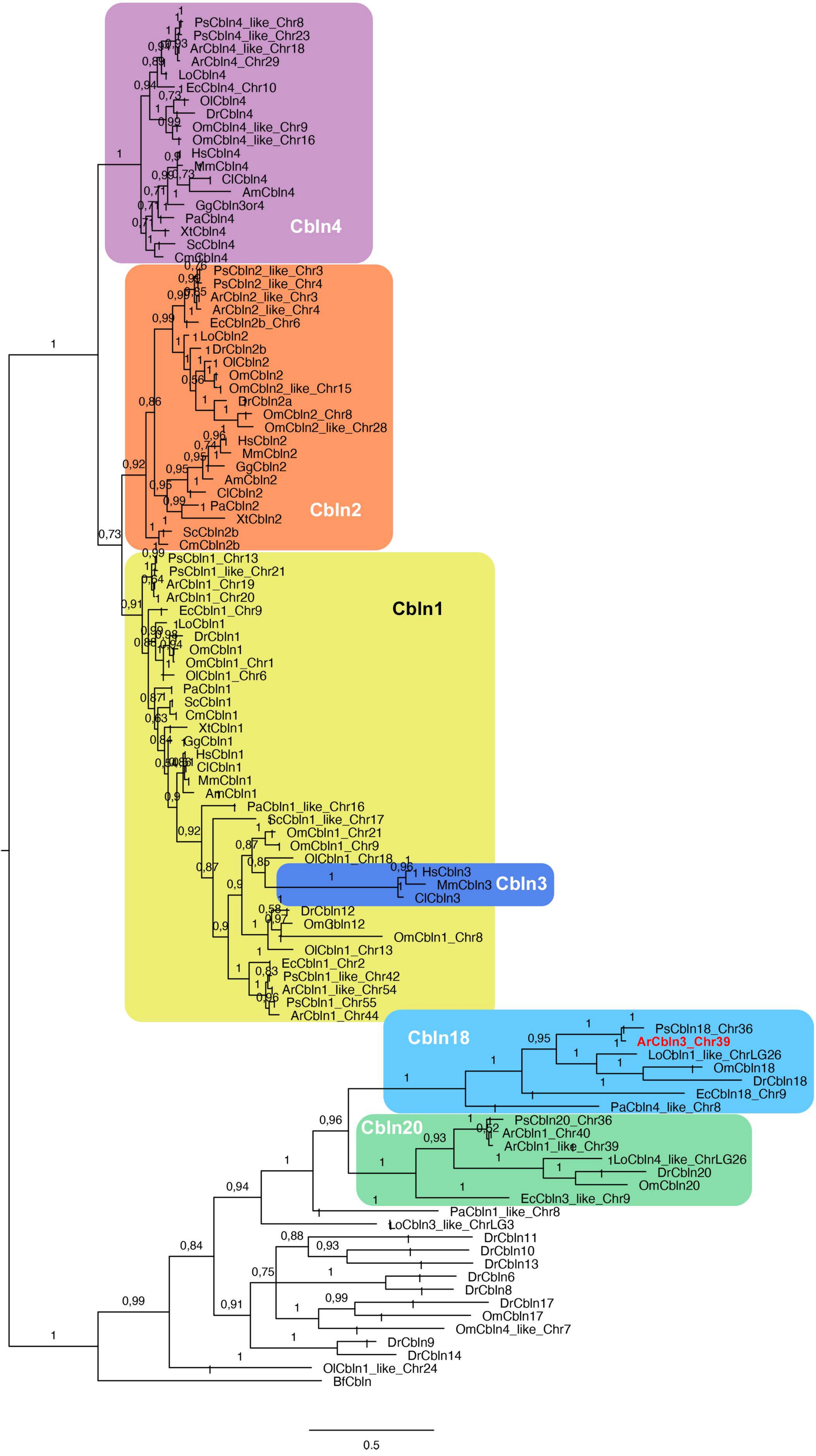


**Supplementary Figure S1: Phylogenetic analysis using MrBayes shows that the sterlet ortholog of the lateral line organ-enriched paddlefish *cerebellin* transcript encodes Cbln18.** Phylogenetic tree of cerebellin family amino acid sequences generated using MrBayes v.3.2.7 (Ronquist et al., 2012). Sequence names reflect the reference-genome annotation and show the chromosomal (Chr) location of the gene if multiple cerebellin genes in that species share the same or similar annotation (e.g., sterlet [Ar] *Cbln1* on chromosomes 19, 20, 40, and 44, and *Cbln1-like* on chromosomes 39 and 54). Maximum support for the Cbln18 clade shows that the lateral line-enriched paddlefish *cerebellin* transcript (Modrell et al., 2017) and its sterlet ortholog (highlighted in bold red font) encode Cbln18 and have been mis-annotated as *Cbln3* in the respective reference genomes (paddlefish GCF_017654505.1; sterlet GCF_902713425.1). Further, the sterlet *Cbln20* ohnologs have been mis-annotated in the reference genome as *Cbln1-like* (chromosome 39) and *Cbln1* (chromosome 40). The tree also shows that Cbln3 (nested within the Cbln1 clade in this tree) is specific to mammals: all non-mammalian sequences annotated as *Cbln3* cluster in other clades. GenBank accession numbers for the sequences used (104 from vertebrates plus the single amphioxus cerebellin sequence) are given in Supplementary Table S2. Species abbreviations: Ar, *Acipenser ruthenus;* Am, *Alligator mississippiensis;* Bf, *Branchiostoma floridae;* Cl, *Canis lupus;* Cm, *Callorhinchus milii;* Dr, *Danio rerio;* Ec, *Erpetoichthys calabaricus;* Gg, *Gallus gallus;* Hs, *Homo sapiens;* Lo, *Lepisosteus oculatus;* Mm, *Mus musculus;* Ol, *Oryzias latipes;* Om, *Oncorhynchus mykiss;* Ps, *Polyodon spathula;* Pa, *Protopterus annectens;* Sc, *Scyliorhinus canicula;* Xt, *Xenopus tropicalis.*

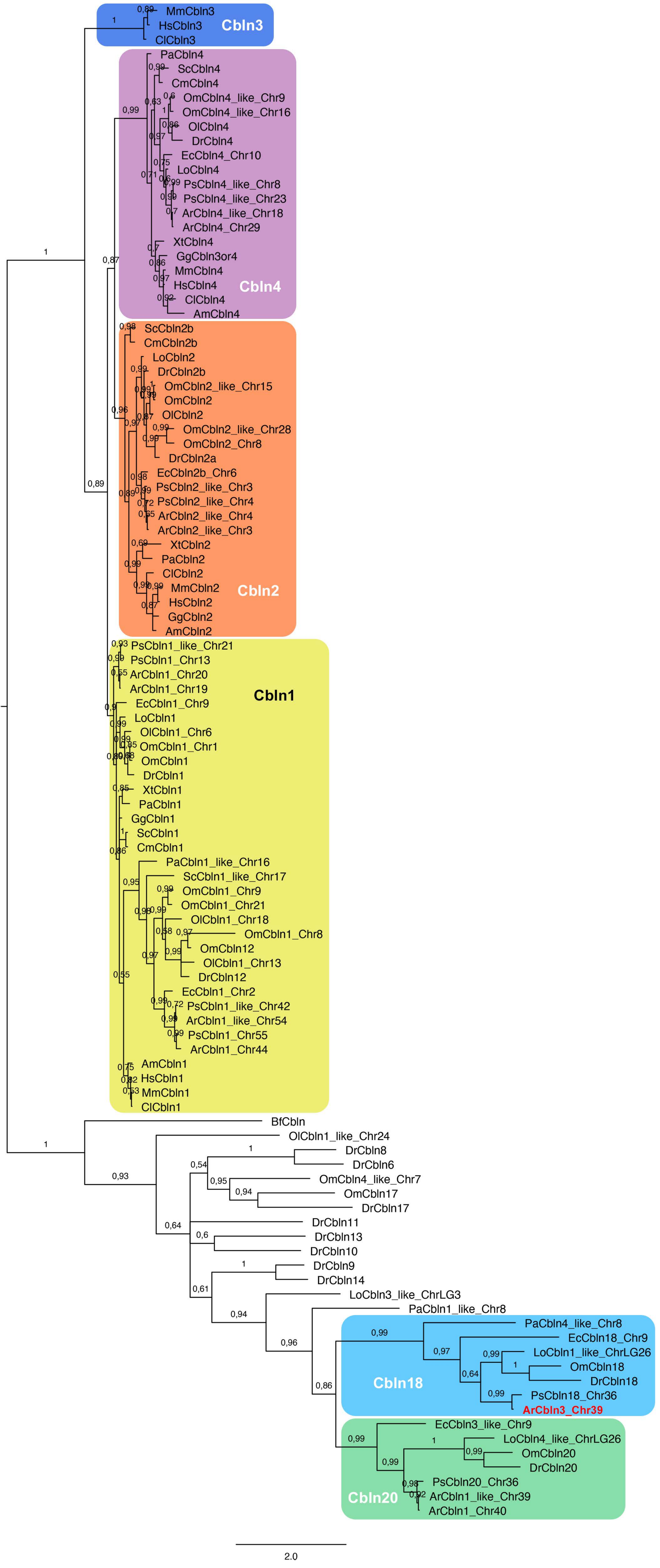


**Supplementary Figure S2: Phylogenetic analysis using PhyloBayes shows that the sterlet ortholog of the lateral line organ-enriched paddlefish *cerebellin* transcript encodes Cbln18.** Phylogenetic tree of cerebellin family amino acid sequences generated using PhyloBayes MPI v1.8c (Lartillot et al., 2013). Sequence names reflect the reference-genome annotation and show the chromosomal (Chr) location of the gene if multiple *cerebellin* genes in that species share the same or similar annotation (e.g., sterlet [Ar] *Cbln1* on chromosomes 19, 20, 40, and 44, and *Cbln1-like* on chromosomes 39 and 54). Maximum support for the Cbln18 clade shows that the lateral line-enriched paddlefish *cerebellin* transcript (Modrell et al., 2017) and its sterlet ortholog (highlighted in bold red font) encode Cbln18 and have been mis-annotated as *Cbln3* in the respective reference genomes (paddlefish GCF_017654505.1; sterlet GCF_902713425.1). Further, the sterlet *Cbln20* ohnologs have been mis-annotated in the reference genome as *Cbln1-like* (chromosome 39) and *Cbln1* (chromosome 40). The tree also shows that Cbln3 is specific to mammals: all non-mammalian sequences annotated as *Cbln3* cluster in other clades. GenBank accession numbers for the sequences used (104 from vertebrates plus the single amphioxus cerebellin sequence) are given in Supplementary Table S2. Species abbreviations: Ar, *Acipenser ruthenus;* Am, *Alligator mississippiensis;* Bf, *Branchiostoma floridae;* Cl, *Canis lupus;* Cm, *Callorhinchus milii;* Dr, *Danio rerio;* Ec, *Erpetoichthys calabaricus;* Gg, *Gallus gallus;* Hs, *Homo sapiens;* Lo, *Lepisosteus oculatus;* Mm, *Mus musculus;* Ol, *Oryzias latipes;* Om, *Oncorhynchus mykiss;* Ps, *Polyodon spathula;* Pa, *Protopterus annectens;* Sc, *Scyliorhinus canicula;* Xt, *Xenopus tropicalis.*
